# A novel induced pluripotent stem cell model of Schwann cell differentiation reveals *NF2*- related gene regulatory networks of the extracellular matrix

**DOI:** 10.1101/2024.05.02.591952

**Authors:** Olivia Lazaro, Sihong Li, William Carter, Oluwamayowa Awosika, Sylvia Robertson, Brooke E. Hickey, Steven P. Angus, Austin House, Wade D. Clapp, Abdul S. Qadir, Travis S. Johnson, Steven D. Rhodes

## Abstract

Schwann cells are vital to development and maintenance of the peripheral nervous system and their dysfunction has been implicated in a range of neurological and neoplastic disorders, including *NF2*-related schwannomatosis. We developed a novel human induced pluripotent stem cell (hiPSC) model to study Schwann cell differentiation in health and disease. We performed transcriptomic, immunofluorescence, and morphological analysis of hiPSC derived Schwann cell precursors (SPCs) and terminally differentiated Schwann cells (SCs) representing distinct stages of development. To validate our findings, we performed integrated, cross-species analyses across multiple external datasets at bulk and single cell resolution. Our hiPSC model of Schwann cell development shared overlapping gene expression signatures with human amniotic mesenchymal stem cell (hAMSCs) derived SCs and *in vivo* mouse models, but also revealed unique features that may reflect species-specific aspects of Schwann cell biology. Moreover, we identified gene co-expression modules that are dynamically regulated during hiPSC to SC differentiation associated with ear and neural development, cell fate determination, the *NF2* gene, and extracellular matrix (ECM) organization. By cross-referencing results between multiple datasets, we identified new genes potentially associated with *NF2* expression. Our hiPSC model further provides a tractable platform for studying Schwann cell development in the context of human disease.

## INTRODUCTION

Schwann cells are glial cells that originate in the neural crest and play a crucial role in myelinating peripheral nerves[1]. Myelination enhances the speed and efficiency of nerve conduction by forming a single spiraling myelin sheath around an axon [2]. Non-myelinating Schwann cells also provide unmyelinated axons with trophic support and mechanical cushioning [3]. Deregulated Schwann cell development can give rise to tumors of the peripheral and cranial nerves called schwannomas, which can have unpredictable clinical behavior, significant morbidities, and exceedingly limited therapeutic options available for their treatment [4].

Vestibular schwannomas are tumors that involve the eight cranial nerves, which are critical for hearing and balance. Affected individuals often suffer significant morbidities including tinnitus, sensorineural hearing loss, and impairment of balance and coordination [5]. Most vestibular schwannomas occur unilaterally; however, approximately 10% of cases involve both sides and are frequently associated with an autosomal dominant, heritable genetic condition called *NF2*- related schwannomatosis (*NF2*-SWN, formally known as neurofibromatosis type 2 [5, 6]. *NF2*- SWN is caused by pathogenic variants in the *NF2* tumor suppressor gene, which encodes Merlin, a cytoskeletal adaptor protein with diverse biochemical functions [7]. The development of bilateral vestibular schwannomas is a hallmark of *NF2*-SWN [8].

To better understand the molecular mechanisms underlying schwannoma genesis in *NF2*-SWN, the development of new model systems for normal and diseased Schwann cells is imperative. Human induced pluripotent stem cells (hiPSCs) have demonstrated promise in modeling a multitude of neurological diseases and cell types, including but not limited to hiPSC-derived astrocytes [9–12], microglia [13–15], and neurons [16, 17]. Direct conversion of human fibroblasts to hiPSC-deviled neurons has also been achieved [17]. These iPSC-derived cells have been employed to interrogate neuropsychiatric diseases [18–20], neurodegenerative diseases [13, 20, 21], and gliomas [22–24]. However, beyond gliomas, considerably less progress has been made in the field of iPSC modeling of peripheral nervous system derived tumors including schwannoma [25].

To address this unmet need, we have developed a novel hiPSC Schwann cell model that can be used to study Schwann cell differentiation in health and disease. Here, we performed transcriptomic profiling of hiPSCs at multiple stages of Schwann cell differentiation to unveil broad networks of differentially expressed genes (DEGs) associated with Schwann cell lineage maturation and compare them to developmental stages outlined in other studies (**Table 1**). To further characterize complex gene regulatory networks (GRNs) associated with Schwann cell differentiation, we employed a systems biology approach, utilizing weighted gene co-expression network analysis (WGCNA) to identify modules of transcriptionally associated genes [26] and to study their temporal dynamics in unison. We further refine the putative roles of these modules using functional enrichment analysis [27]. To validate our findings, we compared the DEGs and gene modules derived from our hiPSC induced Schwann cells with two independent single cell and bulk transcriptomic datasets of human Schwann cell differentiation [28] and mouse sciatic nerve development [29].

**Table 1.**
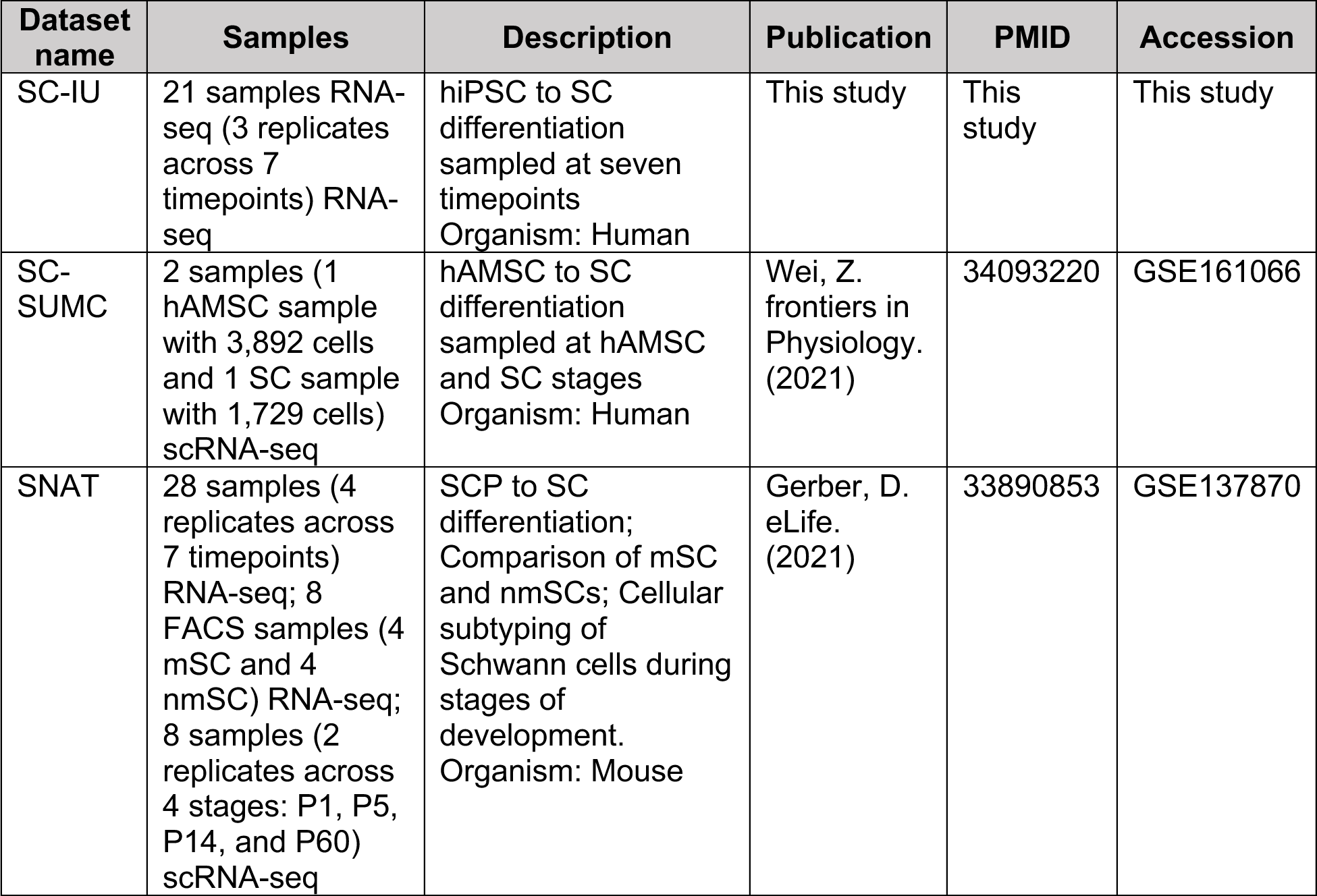
Table of datasets used for analysis in this study.

| Dataset name | Samples | Description | Publication | PMID | Accession |
| --- | --- | --- | --- | --- | --- |
| SC-IU | 21 samples RNA-seq (3 replicates across 7 timepoints) RNA-seq | hiPSC to SC differentiation sampled at seven timepoints<br>Organism: Human | This study | This study | This study |
| SC-SUMC | 2 samples (1 hAMSC sample with 3,892 cells and 1 SC sample with 1,729 cells) scRNA-seq | hAMSC to SC differentiation sampled at hAMSC and SC stages<br>Organism: Human | Wei, Z. *frontiers in Physiology.* (2021) | 34093220 | GSE161066 |
| SNAT | 28 samples (4 replicates across 7 timepoints) RNA-seq; 8 FACS samples (4 mSC and 4 nmSC) RNA-seq; 8 samples (2 replicates across 4 stages: P1, P5, P14, and P60) scRNA-seq | SCP to SC differentiation; Comparison of mSC and nmSCs; Cellular subtyping of Schwann cells during stages of development.<br>Organism: Mouse | Gerber, D. *eLife.* (2021) | 33890853 | GSE137870 |

## METHODS

### hiPSC generation

hiPSC lines were generated from neonatal human dermal fibroblasts (HDFn) from Invitrogen (Catalog #: C-004-5C) using an Epi5™ Episomal iPSC Reprogramming Kit (ThermoFisher Scientific) according to manufacturer’s protocol. Transduced cells were plated on growth factor reduced Matrigel (Corning, 356231) coated culture dishes in the presence of ROCK inhibitor and maintained in Essential 8 (E8) medium. iPSC colonies were mechanically isolated and expanded until at least passage 12 before differentiation. iPSCs were characterized by staining with pluripotency markers and embryoid body (EB) formation. *Mycoplasma* testing was performed prior to Schwann cell differentiation.

### Schwann cell differentiation from hiPSCs

The human iPSC line IBRI.101.J was differentiated into Schwann cells using a six-stage protocol over 31 days as described previously [30]. To start differentiation, the IBRi 101.J hiPSC line was seeded on Matrigel-coated culture plate in E8 media. When the cells reached 50-60% confluency, the culture medium was replaced with Schwann cell basal 1 (SCB1) media (NDM containing 1x N2, 1x B27, 0.005% BSA, 2 mM GlutaMAX, 0.11 mM b-mercaptoethanol in advanced DMEM/F12 and Neurobasal medium (1:1 mix)) with 3 µM CHIR and 20 µM SB431542 for 6 days. On day 7, cells were fed with SCB1 media with 3 µM CHIR, 20 µM SB431542 and 50 ng/mL neuregulin 1 (NRG1) through day 24. SCB1 media was replaced with fresh medium daily, and the cells were routinely dissociated and expanded once they reached 80% confluence with Accutase treatment. On day 25, media was switched to Schwann cell basal 2 (SCB2) media (DMEM, 1x NEAA, 1x BME and 2% FBS) with 200 ng/mL NRG1, 4 ng/mL Forskolin, 100 nM RA-all Trans and 10 ng/mL platelet derive growth factor subunit B (PDGF-BB) for 3 days, then to SCB2 media with 200 ng/mL NRG1 and 10 ng/mL PDGF-BB for 2 days, and finally to SCB2 media with 200 ng/mL NRG1 for 2 more days. By day 31, terminal differentiation to Schwann-like cells was completed. Samples were collected at days 0, 3, 6, 12, 18, 24 and at a uniform 70% confluency in all cases. Cell pellets were collected, and flash frozen at −80°C from three wells for subsequent RNA extraction and RNA-seq, while one well was retained for continued passaging to the next stage.

### Immunoflourescence staining of hiPSC-derived Schwann cell precursors (SCPs) and SCs

We used immunofluorescence (IF) analysis to characterize hiPSC differentiation to SCPs and SCs [31]. For immunostaining, cells were fixed with 4% Paraformaldehyde for 10 minutes. The fixed cells were washed with PBS three times and then blocked and permeabilized with 0.3% Triton X-100 (Sigma) in 1% Bovine Serum Albumin (Life Technologies) and 10% FBS for 1 hour at room temperature. Cells were then incubated with primary antibodies 1:200 dilution of S100β (Abcam #AB52642), 1:50 dilution of p75 (Abcam #AB3125), 1:100 dilution of HOXb7 (Santa Cruz #sc-81292), and 1:500 dilution of GAP43 (Abcam #AB12274) at 4 °C overnight, then with conjugated secondary antibodies at 1:500 dilution for 1 hour at room temperature. The coverslips were mounted on slides with DAPI to stain the nuclei (VectorLabs). Images were captured with an LSM 780 confocal microscope running Zen Black software. Immunocytochemistry analysis confirmed the expression of S100β, p75, HOXb7, and GAP43.

### RNA extraction, library preparation and sequencing

RNA was extracted using phenol-chloroform extraction with TRIzol LS Reagent (Life Technologies Corporation, #10296028). 1mL of TRIzol LS Reagent was added to flash frozen cell pellets followed by 0.2 mL of chloroform. Tubes were shaken by hand for 15 seconds and incubated at room temperature for 2-3 minutes. Centrifugation was performed on samples at 4°C for 15 minutes at 13,200 rpm. The aqueous phase was transferred to an RNase-free tube and an equal amount of 70% ethanol was added to each sample for a final concentration of 35%. Samples were vortexed and binding and washing of RNA was carried out according to the instructions of Invitrogen’s PureLink™ RNA Micro Scale Kit #12183016. Samples were eluted in 20 µL RNase-free water. The quality and quantity of RNA was determined using a NanoDrop™ Spectrophotometer (Thermo Scientific).

A total of 21 samples, 3 replicates for each of the 7 timepoints, were shipped overnight on dry ice to Novogene for quality control, library preparation and RNA-sequencing as further described below.

### Initial processing of RNA-sequencing data

CASAVA (Consensus Assessment of Sequence And Variation) was used for base recognition through the Illumina Genome Analyzer. Raw reads were removed if they contained adapter sequences, greater than 10% undetermined bases, or more than 50% low-quality bases (quality scores less than 5). Sequenced reads were then aligned to the GRCh38 Genome using HISAT2 v2.0.5 and quantified with feature counts v1.5.0-p3. FPKM values were calculated from the read counts and CDS lengths for each gene across all timepoints and replicates.

### Differential Gene Expression analysis

Differential gene expression analysis was performed with edgeR v3.22.5. The log2 fold change (Log2FC) was calculated between groups and the p-values (P) were corrected using the Benjamini & Hochberg method (BH) to account for the false discovery rate (FDR). Each timepoint (day 3, 6, 12, 18, 24, 31) was compared to baseline (day 0), resulting in 6 pairwise comparisons. DEGs with Log2FC > 5.0 and FDR < 0.001 were identified for each comparison. Functional enrichment analysis was performed on DEGs from each comparison using clusterProfiler v3.8.1 to test the statistical enrichment for genes within the Reactome, DO (Disease Ontology), and DisGeNET databases. Terms with FDR < 0.05 were considered enriched. Functional enrichment on other gene sets derived from this analysis was performed using ToppGene [32].

### Network Construction and Module Detection with WGCNA and lmQCM

Statistical analysis was performed using R (v4.1.1.). Genes with the lowest 10% mean FPKM across all timepoints were removed. Ensembl IDs for the filtered genes were converted to HGNC gene symbols using biomaRt (v2.52.0) and the Ensembl gene IDs from the Genome assembly GRCh38.p14. For HGNC symbols with multiple corresponding Ensembl IDs, the Ensembl ID with the highest average FPKM was used. WGCNA module generation was conducted with an unsigned Pearson correlation coefficient (PCC) matrix. Soft thresholding power was calculated to reduce noise for gene correlations and emphasize correlations between genes per WGCNA documentation. A block size of 22000 was used for network construction and module detection as described previously [26]. Eigengenes were subsequently calculated for each module detected by WGCNA. Gene modules whose eigengene values changed significantly (P<0.05) for at least one timepoint were identified by ANOVA. Enrichment analysis was performed using clusterProfiler to identify significantly associated ontology terms for each module. All ontologies contained in clusterProfiler were evaluated, including BP (biological process), CC (cellular component), and MF (molecular function).

### Evaluating Schwann cell differentiation from amniotic mesenchymal stem cells via single cell RNA sequencing

Single cell RNA-seq (scRNA-seq) data of Schwann cells (SCs) differentiated from human amniotic mesenchymal stem cells (hAMSCs) was downloaded from Gene Expression Omnibus (GSE161066) [28]. Seurat [33] was used for initial data cleaning and quality control, removing cells with >2,500 or <200 unique feature counts as well as cells with >5% mitochondrial RNAs. The hAMSC and SCs samples were each normalized with SCTransform [34, 35] and then integrated with Harmony [36, 37]. Uniform manifold approximation and projection (UMAP) [38] was used as a final dimensionality reduction step. The integrated object was clustered, markers were identified for each cluster (|Log2FC|>0.58 and *FDR<0.05*), and the relative cluster proportions were compared between hAMSCs and SCs. DGE analysis was also performed between hAMSC and SCs to identify DEGs specific to either group (|Log2FC|>0.58 and *FDR<0.05*).

### Evaluation of Schwann cell differentiation in mouse sciatic nerve

Bulk RNAseq data (FPKM values) from sciatic nerve harvested at various developmental stages (E17.5, P1, P5, P14, P24, P60) and peripheral nerves distal to the DRG at E13.5 were downloaded from the SNAT data portal/GEO (GSE137870) [29]. For gene names with more than one ID, the ID was selected with the highest average expression across all cells in the dataset. Because gene to gene comparisons are biases by sequence length, we converted the FPKM values to TPM values for gene filtering. Genes with a TPM value greater than the median TPM value of all genes were retained for downstream analysis. Next, differential gene expression analysis was performed using edgeR (3.34.1). Six comparisons were performed, one for each timepoint (E17.5, P1, P5, P14, P24, P60) compared to the E13.5 reference timepoint. Genes were considered differentially expressed if they had an |Log2FC|>5 and an *FDR<0.001*. Genes were mapped to their appropriate human homolog using biomaRt (2.48.3) and the Dec 2021 Ensembl archive for both human and mouse.

### Evaluation of myelinating compared to non-myelinating mouse Schwann cells

Bulk RNAseq data (FPKM values) from sorted Schwann cell populations enriched for myelinating versus non-myelinating Schwann cells were downloaded from the SNAT data portal/GEO (GSE137947) [29]. The initial processing and homolog mapping steps were as described above. DEGs were compared between myelinating and non-myelinating Schwann cells. Genes were considered differentially expressed if they had an |Log2FC|>5 and an *FDR<0.001*.

### Cellular subtyping of Schwann cells from mouse sciatic nerve

The integrated Seurat (4.3.0) object from the SNAT data portal (SS2_all) was downloaded [29] for analysis and comparison with additional datasets. UMAP was calculated on the integrated components using the first 10 dimensions. Marker genes were identified using the FindAllMarkers function. The homolog mapping steps were as previously described. In addition to identifying markers from the previously defined clusters, we also determined DEGs for various differentiation trajectories. Specifically, we identified DEGs using the FindMarkers function contrasting immature Schwann cells (iSCs) from myelinating Schwann cells (mSCs), i.e., iSC vs. tSC and nm(R)SC, and DEGs contrasting iSCs from non-myelinating Schwann cells (nmSCs), i.e., iSC vs. pmSC, mSC cluster 1, mSC cluster 2, and mSC cluster 3. Genes were considered differentially expressed if they had an |Log2FC|>0.58 and an *FDR<0.05*.

### Identification of conserved gene expression programs in differentiating Schwann cells across myelination status and species

To examine the similarity of transcriptional programs between mouse and hiPSCs during Schwann cell differentiation, we compared DEGs from our SC-IU dataset with the SNAT RNA-seq dataset for each of the six time point comparisons. The Jaccard index (JI) was calculated for up-regulated DEGs for each pairwise comparison between the SC-IU and SNAT datasets (i.e., E17.5 up-regulated DEGs (SNAT) and Day 3 up-regulated DEGs (SC-IU) JI, E17.5 up-regulated DEGs (SNAT) and Day 6 up-regulated DEGs (SC-IU) JI, etc.). The same process was conducted for the down-regulated DEGs, resulting in a total of 36 total JI values for the up-regulated and downregulated DEGs, respectively. We then averaged and normalized the JI values to obtain a square matrix with row and column sums of 1 to account for any skew toward either end of the differentiation process. We then repeated this process with less stringent DEG calling criteria (|Log2FC|>0.58 and an *FDR<0.05*). The consistency of the JI matrix was evaluated using a Student’s t-test of the diagonal of the matrix vs. the off-diagonal and a Spearman correlation of the JI with the distance of each cell from the diagonal. Next, we compared the union of up-regulated and down-regulated DEGs from both the SC-IU and SNAT datasets to differentiation and developmental gene modules from the WGCNA analysis using the original criteria (|Log2FC|>5 and an *FDR<0.001*). Finally, mSC markers and nmSC markers from the SNAT scRNA-seq dataset were compared to the mSC DEGs identified from the SNAT myelination bulk RNA-seq dataset. We use these comparisons to evaluate the consistency of genes across modalities, species, myelination status, and analysis technique to identify high-confidence targets for the study of Schwann cell differentiation.

## RESULTS

### Morphological and immunofluorescence-based validation of hiPSC induced differentiation to SPCs and SCs

To confirm robust differentiation of hiPSCs into Schwann cells precursors (SCPs) and subsequently SCs, we performed morphological and immunofluorescence (IF) staining of cells collected at seven timepoints along the hiPSC to SC continuum (**Fig. 1A**, **Table 1**). We observed morphological changes during the differentiation process where the cells exhibited increasing levels of cohesion into aggregates and reduced levels of random arrangements of individual cells (**Fig. 1A**). Schwann cells are derived from the neural crest and characterized by the expression of distinct makers including S100β, p75, HOXb7, and GAP43 (**Fig. 1B**). A gradual increase in the expression of S100β, p75, HOXb7, and GAP43 was observed throughout the differentiation process reaching a maximum at the SC stage (**Fig. 1B, Fig S1**). Collectively, these findings indicate that our methodology can efficiently generate SCPs and SLPs from hiPSC with high fidelity.

**Fig. 1.**
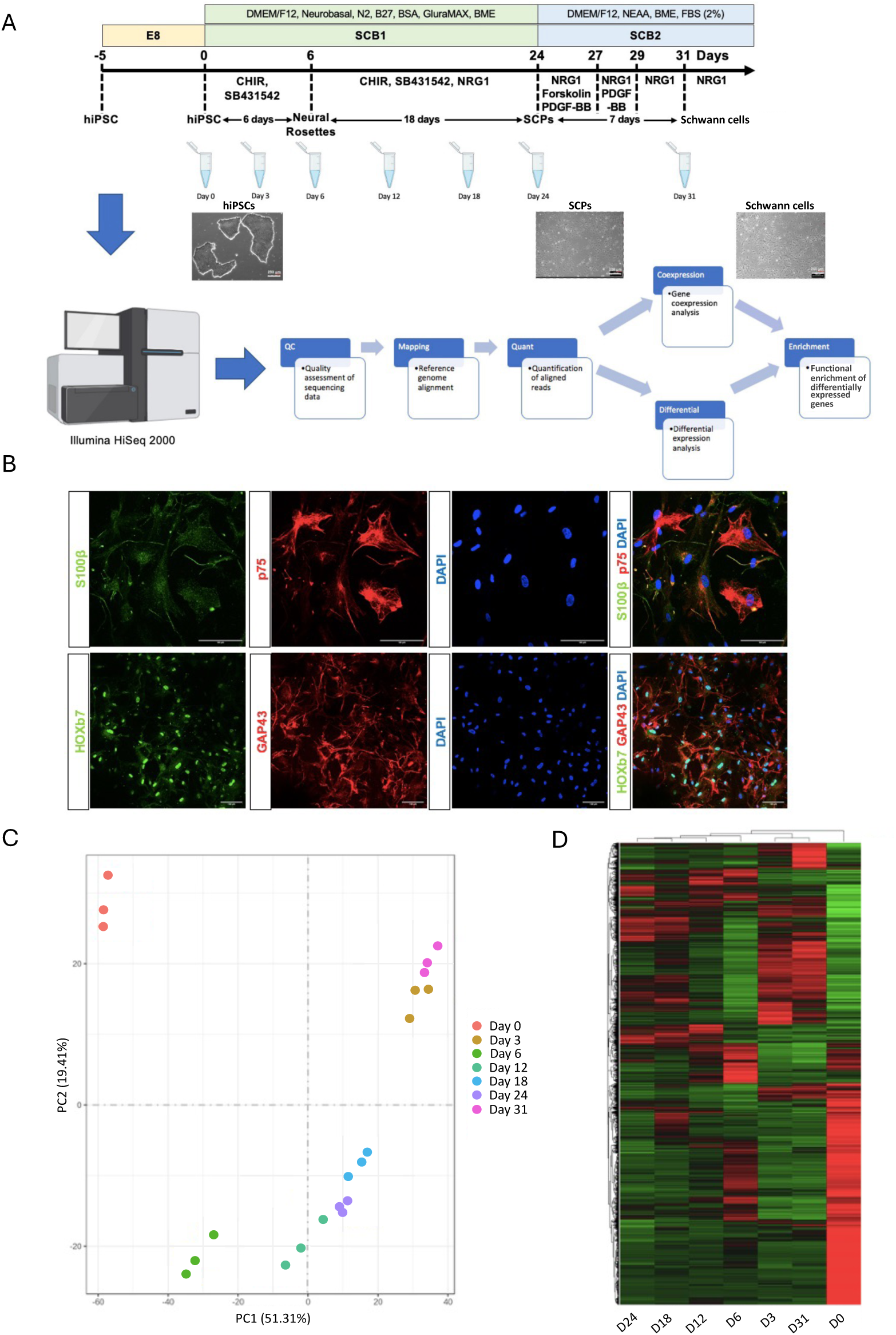
Workflow and validation of directed differentiation of hiPSCs into human Schwann cells. **A)** Schematic representation of the differentiation of hiPSCs into Schwann cell precursors (SCPs) and Schwann cells (SCs) (Upper panel). Representative bright-field images showing the process of differentiation into SCPs and SCs. Scale bar, 100 µm (middle panel). Steps involve sequencing and data analysis (lower panel). **B)** Schwann cell differentiation was confirmed by positive immunocytochemical staining for S100β (green), p75 (red) in upper panel and HOXb7 (green), GAP43 (red) in lower panel after 31 days of differentiation. DAPI (blue) was used to stain the cell nuclei. Scale bars, 100 µm. **C)** Timepoints can be stratified into stages of differentiation by PCA. **D)** Heatmap of differentially expressed genes across different timepoints where the rows (genes) were clustered using hierarchical clustering.

### Differential gene expression during SC differentiation

To characterize gene expression dynamics during Schwann cell differentiation, we performed RNA sequencing of cells collected at seven discrete stages along the hiPSC to SC continuum. As anticipated, replicates from each stage of differentiation clustered together in PCA space (**Fig. 1C**). Notably, the hiPSC stage had clear separation from all other stages (**Fig. 1C**) and the neural rosette (NR) through SCP stages of differentiation clustered more closely together compared to other stages (**Fig. 1C**). There were considerable transcriptional differences between all stages of differentiation, most notably hiPSCs versus later stages of differentiation (**Fig. 1D, Fig. S1-S3**). We identified 18 genes that were significantly up-regulated across all 6 comparisons (**Table 2**). Of these, *ANXA1* and *CDH6* were also among the top 5 most significant up-regulated DEGs (lowest FDR adjusted p-values) within any of the timepoint comparisons (**Table 2**). Notably, *CDH6* is associated with cell-cell adhesion and *ANXA1* has been implicated in Schwann cell proliferation and migration [47]. Of the 30 total top 5 up-regulated DEGs per timepoint comparison, 18 were up-regulated across all timepoints These 18 genes were enriched for ontology terms related to cell adhesion (GO:0007155, *P<0.001*) and anatomical structure formation involved in morphogenesis (GO:0048646, *P<0.001*). In total, 50 significantly down-regulated DEGs were identified at each timepoint comparison (**Table 2**). However, only four were also found in the top 5 most significant down-regulated DEGs (lowest FDR adjusted p-values) at any of the timepoint comparisons (**Table 2**). Of the top 5 down-regulated DEGs per timepoint comparison, 18 genes were down-regulated across all timepoints and were enriched for ontology terms related to the regulation of cell fate specification (GO:0042659, *P<0.001*).

**Table 2.** Top 5 up- and down-regulated DEGs for each of the 6 timepoints (k) (days 3, 6, 12, 18, 24, and 31) compared to day 0 hiPSCs as the reference.

| Day k vs. Day 0 | Gene | Log2FC | P | FDR |
| --- | --- | --- | --- | --- |
| Day 3 Up | *COL8A1* | 11.33 | 0 | 0 |
| Day 3 Up | *POSTN* | 10.15 | 0 | 0 |
| Day 3 Up | *SERPINE1* | 9.22 | 0 | 0 |
| Day 3 Up | *IGFBP3* | 7.61 | 0 | 0 |
| Day 3 Up | *TGFBI* | 7.15 | 0 | 0 |
| Day 3 Down | *CD24* | -6.37 | 5.14E-260 | 8.69E-257 |
| Day 3 Down | *EPCAM* | -7.93 | 1.19E-229 | 1.07E-226 |
| Day 3 Down | *HLA-A* | -5.53 | 5.65E-213 | 4.16E-210 |
| Day 3 Down | *PMEL* | -7.55 | 7.08E-213 | 5.08E-210 |
| Day 3 Down | *ESRG* | -9.76 | 7.22E-212 | 5.06E-209 |
| Day 6 Up | *HAPLN1* | 6.97 | 8.78E-236 | 2.53E-231 |
| Day 6 Up | *CDH6* | 5.91 | 1.68E-227 | 2.43E-223 |
| Day 6 Up | *PRTG* | 4.91 | 4.72E-225 | 4.54E-221 |
| Day 6 Up | *WLS* | 5.36 | 1.57E-184 | 7.55E-181 |
| Day 6 Up | *GREB1L* | 6.90 | 2.12E-183 | 8.73E-180 |
| Day 6 Down | *TDGF1* | -5.01 | 8.98E-198 | 6.47E-194 |
| Day 6 Down | *ADCY2* | -6.77 | 1.89E-190 | 1.09E-186 |
| Day 6 Down | *LINC00678* | -5.15 | 2.72E-151 | 7.83E-148 |
| Day 6 Down | *AC064802.1* | -4.65 | 9.03E-148 | 2.37E-144 |
| Day 6 Down | *DUSP6* | -4.65 | 7.95E-145 | 1.64E-141 |
| Day 12 Up | *ANXA1* | 10.95 | 4.31E-276 | 6.09E-272 |
| Day 12 Up | *CPE* | 5.72 | 2.35E-191 | 1.67E-187 |
| Day 12 Up | *PLK2* | 5.42 | 1.95E-190 | 1.11E-186 |
| Day 12 Up | *PRTG* | 4.52 | 1.72E-185 | 8.13E-182 |
| Day 12 Up | *CDH6* | 5.53 | 4.22E-181 | 1.71E-177 |
| Day 12 Down | *TDGF1* | -8.16 | 0 | 0 |
| Day 12 Down | *AC064802.1* | -8.52 | 7.05E-207 | 6.65E-203 |
| Day 12 Down | *NANOG* | -7.30 | 7.96E-173 | 2.82E-169 |
| Day 12 Down | *USP44* | -5.13 | 5.67E-164 | 1.46E-160 |
| Day 12 Down | *AC009446.1* | -7.39 | 2.73E-162 | 6.42E-159 |
| Day 18 Up | *ANXA1* | 10.79 | 0 | 0 |
| Day 18 Up | *COL8A1* | 10.92 | 0 | 0 |
| Day 18 Up | *CAV1* | 6.12 | 0 | 0 |
| Day 18 Up | *SERPINE1* | 8.14 | 5.48E-295 | 3.85E-291 |
| Day 18 Up | *POSTN* | 8.01 | 2.79E-294 | 1.56E-290 |
| Day 18 Down | *TDGF1* | -10.73 | 2.01E-273 | 7.06E-270 |
| Day 18 Down | *USP44* | -5.52 | 1.62E-253 | 4.54E-250 |
| Day 18 Down | *ZSCAN10* | -6.42 | 3.59E-216 | 4.03E-213 |
| Day 18 Down | *ESRG* | -5.16 | 8.92E-199 | 7.82E-196 |
| Day 18 Down | *VSNL1* | -7.28 | 3.70E-179 | 2.16E-176 |
| Day 24 Up | *ANXA1* | 11.19 | 0 | 0 |
| Day 24 Up | *POSTN* | 8.43 | 0 | 0 |
| Day 24 Up | *CAV1* | 6.16 | 0 | 0 |
| Day 24 Up | *PLK2* | 5.77 | 0 | 0 |
| Day 24 Up | *SPP1* | 4.22 | 1.63E-291 | 9.33E-288 |
| Day 24 Down | *USP44* | -5.49 | 8.04E-274 | 2.29E-270 |
| Day 24 Down | *ESRG* | -5.53 | 8.55E-249 | 1.22E-245 |
| Day 24 Down | *ZSCAN10* | -7.03 | 1.28E-236 | 1.52E-233 |
| Day 24 Down | *PMEL* | -4.70 | 6.92E-211 | 6.17E-208 |
| Day 24 Down | *B3GNT7* | -6.79 | 7.86E-201 | 5.98E-198 |
| Day 31 Up | *CTGF* | 6.05 | 0 | 0 |
| Day 31 Up | *VIM* | 6.55 | 0 | 0 |
| Day 31 Up | *COL5A2* | 7.04 | 0 | 0 |
| Day 31 Up | *COL8A1* | 11.75 | 0 | 0 |
| Day 31 Up | *POSTN* | 9.95 | 0 | 0 |
| Day 31 Down | *CD24* | -6.94 | 0 | 0 |
| Day 31 Down | *HLA-A* | -6.02 | 8.07E-296 | 7.22E-293 |
| Day 31 Down | *SFRP2* | -5.66 | 3.19E-289 | 2.61E-286 |
| Day 31 Down | *L1TD1* | -5.78 | 4.22E-275 | 3.26E-272 |
| Day 31 Down | *EPCAM* | -7.92 | 1.74E-267 | 1.18E-264 |

### Gene co-expression modules associated with differentiation, axon guidance and ECM

Building on the results of our differential gene expression analysis, we next performed weighted gene co-expression network analysis (WGCNA) to identify co-regulated gene modules during hiPSC differentiation to SCs. WGCNA identified 24 modules (**Fig. 2A**), and 11 of these had significantly altered eigengene values during differentiation (i.e., *P<0.05*, **Table 3**). Of the 11 differentiation-associated modules, 10 had significantly enriched ontology terms associated with them (**Table 3**). The most significant module identified by ANOVA was the brown module (*P<0.001*, **Table 3**, **Fig. 2B**) which had an eigengene peak at day 6, i.e., a significant increase from day 3 to day 6 (*P<0.001*, **Fig. 2B**) followed by a decrease from day 6 to day 12 (*P<0.001*, **Fig. 2B**). Given the close relationship between Schwann cell development and neurons, we were intrigued that the brown module was functionally enriched for axon guidance (P<0.001, **Fig. 2C**), forebrain development (*P<0.001*, **Fig. 2C**), and sensory organ morphogenesis (*P<0.001*, **Fig. 2C**).

**Fig. 2.**
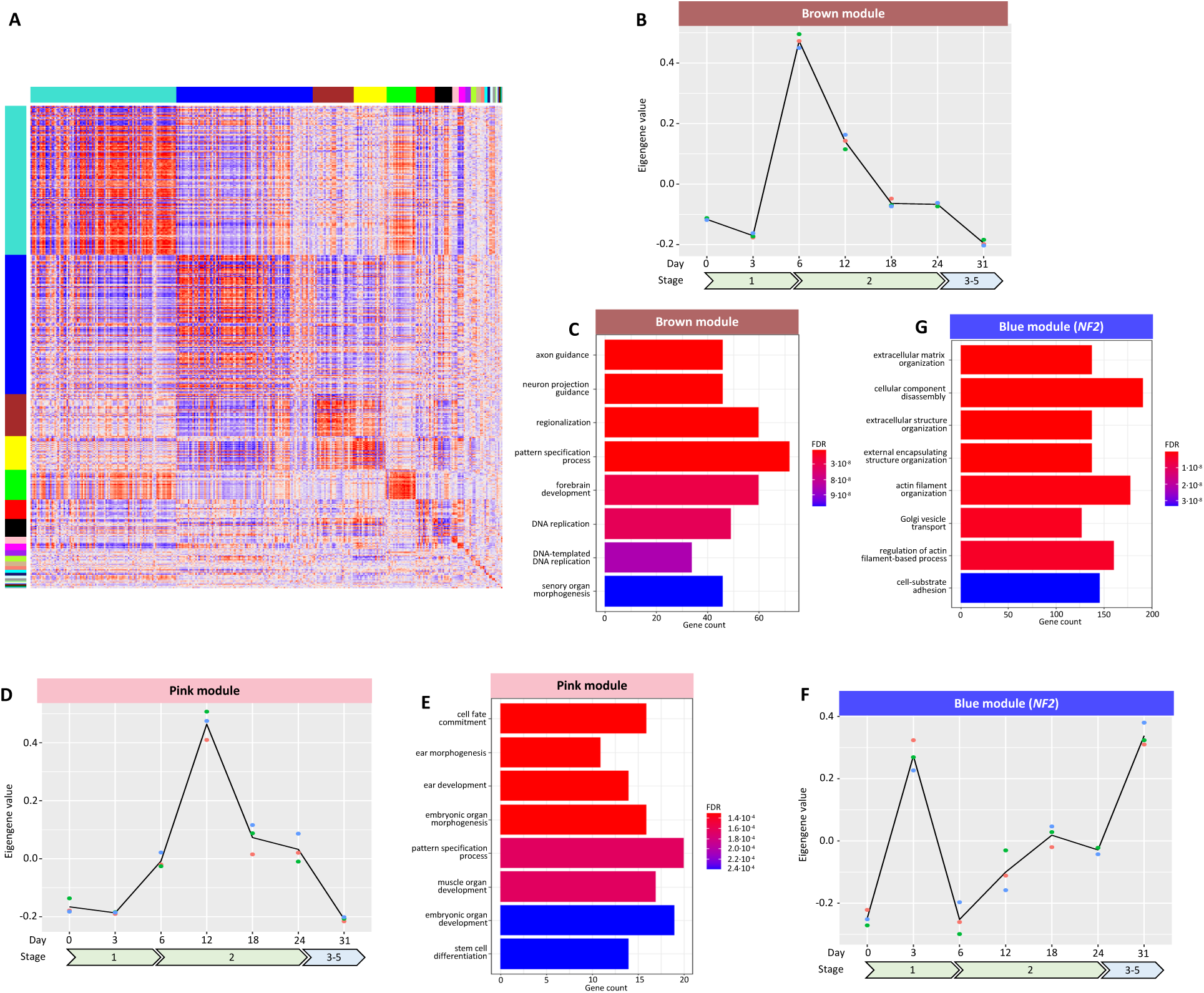
WGCNA reveals dynamic changes in module eigengene values during hiPSC to SC differentiation related to tissue structure and neural development. **A)** Gene correlation matrix with different modules annotated. **B)** Brown module eigengene values are plotted over the course of the study timepoints. **C)** Significant ontology terms from functional enrichment analysis for the brown module. **D)** Pink module eigengene changes over the study timepoints. **E)** Significant ontology terms from functional enrichment analysis for the pink module. **F)** Blue module (containing *NF2*) eigengene changes over the study timepoints. **G)** Significant ontology terms from functional enrichment analysis for the blue module (containing *NF2*).

**Table 3.** Significantly altered gene modules in relation to timepoint evaluated by ANOVA. The number of genes in the module, ANOVA P, and most significant GO term are listed for each module. Modules with no significant GO terms are blank.

| Module | Genes | P | Top GO term | File |
| --- | --- | --- | --- | --- |
| Brown | 1449 | 7.66E-17 | Axon guidance | File S2 |
| Red | 845 | 2.63E-14 | Organic hydroxy compound biosynthetic process | File S3 |
| Turquoise | 4615 | 5.23E-14 | Nuclear chromosome segregation | File S4 |
| Cyan | 100 | 1.38E-11 |  | File S5 |
| Pink | 692 | 2.11E-11 | Cell fate commitment | File S6 |
| Blue | 3797 | 2.16E-10 | Extracellular matrix organization | File S7 |
| Black | 833 | 1.73E-8 | Intraciliary transport particle B | File S8 |
| Magenta | 159 | 4.99E-5 | Endoplasmic reticulum protein-containing complex | File S9 |
| Yellow | 1288 | 1.52E-2 | Cytoplasmic translation | File S10 |
| Midnight Blue | 78 | 4.19E-2 | Transcription coregulator activity | File S11 |
| Green Yellow | 146 | 4.26E-2 | Methyltransferase activity | File S12 |

The next two most significant modules were red (*P<0.001*) and turquoise (*P<0.001*) (**Table 3**). The red module eigengene steadily increased from hiPSC to SCP stage (*P<0.001*, i.e., day 0 to day 24, **Fig. S4**) then dramatically decreased from SCP to SC stage (*P<0.001*, i.e., day 24 to day 31, **Fig. S4**) and was functionally enriched for biosynthetic pathways (**Fig. S4**). In contrast, the turquoise module eigengene, which was functionally enriched for functionally enriched for exocytosis (*P<0.001*, **Fig. S5**) and regulation of neuron projection development (*P<0.001*, **Fig. S5**) decreased dramatically from the hiPSC to preneural rosette (pre-NR) stage (*P<0.001*, i.e., day 0 to day 3, **Fig. S5**) and remained suppressed through terminal SC differentiation (*P=0.732*, i.e., day 3 to day 31, **Fig. S5**). Another pair of modules that showed distinct expression patterns were the pink and blue modules (*P<0.001* and *P<0.001* respectively, **Table 3**). Similar to the brown module that had a distinct eigengene peak at day 6, the pink module peaked at day 12, i.e., a significant increase from day 6 to day 12 (*P<0.001*, **Fig. 2D**) and decreased thereafter from day 12 to day 18 (*P<0.001*, **Fig. 2D**). Not surprisingly, this module had numerous ontology terms related to cell fate commitment and ear development (**Fig. 2E**). The blue module eigengene also had a peak early in hiPSC to SC differentiation at day 3, i.e., a significant increase from day 0 to day 3 (*P<0.001*, **Fig. 2F**) and decrease from day 3 to day 6 (*P<0.001*, **Fig. 2F**). However, this module eigengene gradually increased again from NR through completion of SC maturation (*P<0.001*, **Fig. 2F**). Notably, the blue module contains the *NF2* tumor suppressor gene and had significant functional enrichment for extracellular matrix organization (*P<0.001*, **Fig. 2G**), extracellular structure organization (*P<0.001*, **Fig. 2G**), and cell-substrate adhesion (*P<0.001*, **Fig. 2G**). Taken together with the remaining modules (**Fig. S6-S11**), the results of our WGCNA reveal that hiPSC differentiation to SCs is associated with dynamic changes in gene expression that are reflective of Schwann cell biological function and show associations with neural morphology, ear development, and cellular organization.

### Signatures of hAMSC to SC differentiation are unique at the single cell level

To evaluate the robustness and generalizability of our findings, we extended our analysis to examine whether similar gene modules and pathways were retained in other complementary models of human Schwann cell differentiation. Wei and colleagues performed single cell RNA-sequencing of SCs derived from human amniotic mesenchymal stem cells (hAMSCs) to identify heterogenous SC populations [28] (SC-SUMC: GSE161066, **Table 1**). The scRNA-seq data contained 3,892 hAMSCs and 1,729 SCs. After quality control (QC) and dimensionality reduction, we identified a total of 16 clusters (**Fig. 3A**). DGE analysis revealed clear differences between hAMSC and SCs. Some of the most upregulated genes in SC compared to hAMSCs included ECM related genes like: *COL1A1* and *COL3A1* (**Fig. 3B**, **Table 4**). To further characterize each cluster, we performed cluster-specific marker analysis using Seurat to identify distinct gene sets highly expressed in each cluster (**Fig. 3C**). We observed that some clusters had a higher proportion of either hAMSCs or SCs, suggesting that they may represent different stages of differentiation **(Fig. 3D**). For example, cluster 8 and 14 comprised of a higher proportion of SC sample compared to the hAMSC sample (**Fig. 3D**) and both of these clusters overexpressed ECM-related markers. Cluster 8 overexpressed *VIM*, and cluster 14 overexpressed *ITGA1*, *COL1A1*, and *ADAMTS6* (**Fig. 3C**, **Table S1**).

**Fig. 3.**
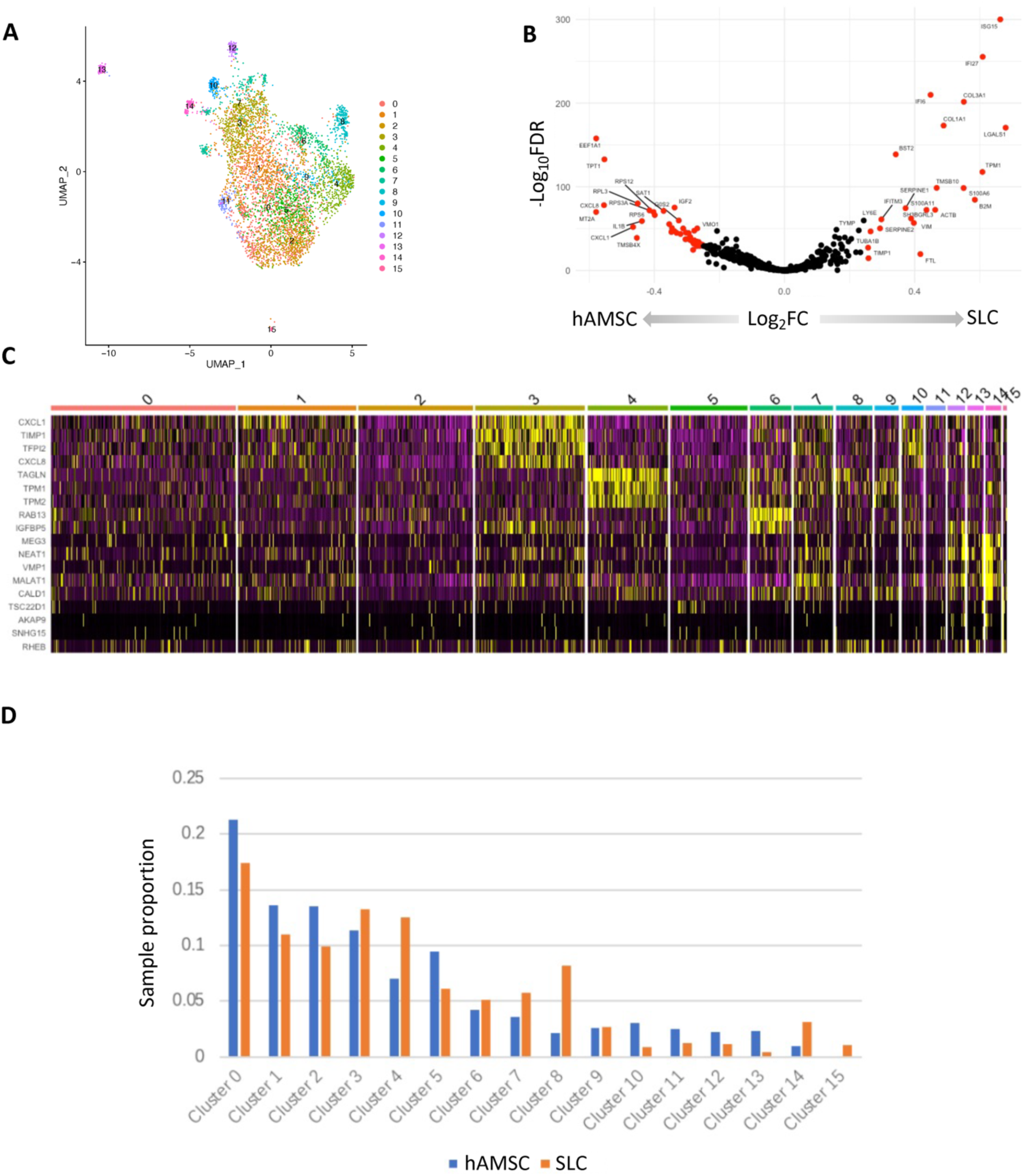
SC-SUMC scRNA-seq analysis. **A)** UMAP clustering of hAMSCs and SCs from the integrated SC-SUMC dataset (GSE161066) reveals 16 clusters. **B)** Volcano plot of differentially expressed genes when compared between hAMSCs and SCs. **C)** Heatmap of cluster specific markers. **D)** Proportion of hAMSCs and SCs comprising each cluster from the integrated SC- SUMC dataset.

**Table 4.** Top 10 up- and down-regulated DEGs in hAMSCs versus SCs from the SC-SUMC dataset.

| Gene | Proportion hAMSCs>0 | Proportion SCs>0 | Log<sub>2</sub>FC | P | FDR |
| --- | --- | --- | --- | --- | --- |
| *ISG15* | 0.04 | 0.44 | 0.66 | 1.00E-300 | 0.00E+00 |
| *IFI27* | 0.02 | 0.32 | 0.61 | 4.98E-256 | 5.71E-252 |
| *IFI6* | 0.01 | 0.27 | 0.45 | 1.37E-210 | 1.58E-206 |
| *COL3A1* | 0.07 | 0.40 | 0.55 | 2.62E-202 | 3.00E-198 |
| *COL1A1* | 0.06 | 0.34 | 0.49 | 6.85E-174 | 7.86E-170 |
| *LGALS1* | 0.49 | 0.81 | 0.68 | 2.36E-171 | 2.71E-167 |
| *EEF1A1* | 0.97 | 0.88 | -0.58 | 1.66E-158 | 1.90E-154 |
| *BST2* | 0.02 | 0.22 | 0.34 | 1.86E-139 | 2.14E-135 |
| *TPT1* | 0.96 | 0.88 | -0.55 | 1.79E-133 | 2.05E-129 |
| *TPM1* | 0.31 | 0.60 | 0.61 | 1.85E-118 | 2.12E-114 |
| *TMSB10* | 0.88 | 0.95 | 0.47 | 2.17E-99 | 2.49E-95 |
| *S100A6* | 0.47 | 0.70 | 0.55 | 3.15E-99 | 3.62E-95 |
| *RPL3* | 0.83 | 0.66 | -0.45 | 8.45E-81 | 9.70E-77 |
| *CXCL8* | 0.71 | 0.48 | -0.55 | 9.51E-79 | 1.09E-74 |
| *IGF2* | 0.20 | 0.01 | -0.34 | 6.67E-76 | 7.65E-72 |
| *RPS3A* | 0.80 | 0.61 | -0.42 | 1.52E-72 | 1.75E-68 |
| *SAT1* | 0.35 | 0.12 | -0.37 | 7.64E-72 | 8.77E-68 |
| *MT2A* | 0.98 | 0.98 | -0.58 | 1.37E-70 | 1.57E-66 |
| *RPS12* | 0.89 | 0.77 | -0.40 | 1.54E-70 | 1.77E-66 |
| *RPS6* | 0.76 | 0.57 | -0.40 | 6.27E-67 | 7.20E-63 |

### Evaluation of Schwann cell differentiation in mouse sciatic nerve

To further validate our hiPSC model of Schwann cell differentiation *in vitro*, we further compared our transcriptomic data with an *in vivo* dataset comprehensively cataloging Schwann cell development in the mouse sciatic nerve. Gerber and colleagues generated gene expression profiles from Schwann cells isolated from mouse sciatic nerve throughout embryogenesis and postnatal development, which are publicly available through the Sciatic Nerve Atlas, i.e., SNAT, [29]. By integrating bulk and single cell RNA-seq data from SNAT, we identified unique and conserved gene signatures of Schwann cell differentiation across species and modalities.

In our SC-IU dataset, a total of 50 genes were significantly up-regulated across all timepoint comparisons, i.e., were contained in each comparison (**Fig. 4A-F, Table S2, Table S3**). These genes represented myelination (GO:0042552, *P<0.001*) and collagen-containing extracellular matrix (GO:0062023, *P<0.001*). The 362 down-regulated DEGs significant at all comparisons were enriched for neuron development (GO:0048666, *P<0.001*) and synaptic signaling (GO:0099536, *P<0.001*). Furthermore, we identified 9 gene co-expression modules that were significantly altered over developmental stage from E13.5 to P60. (**Table S4**, **Fig. S12-S21**). Notably some of these modules were also enriched for terms related to neural development like the SNAT-brown module (GO:0010975, Regulation of neuron projection development, *P<0.001,* **Table S4**).

**Fig. 4.**
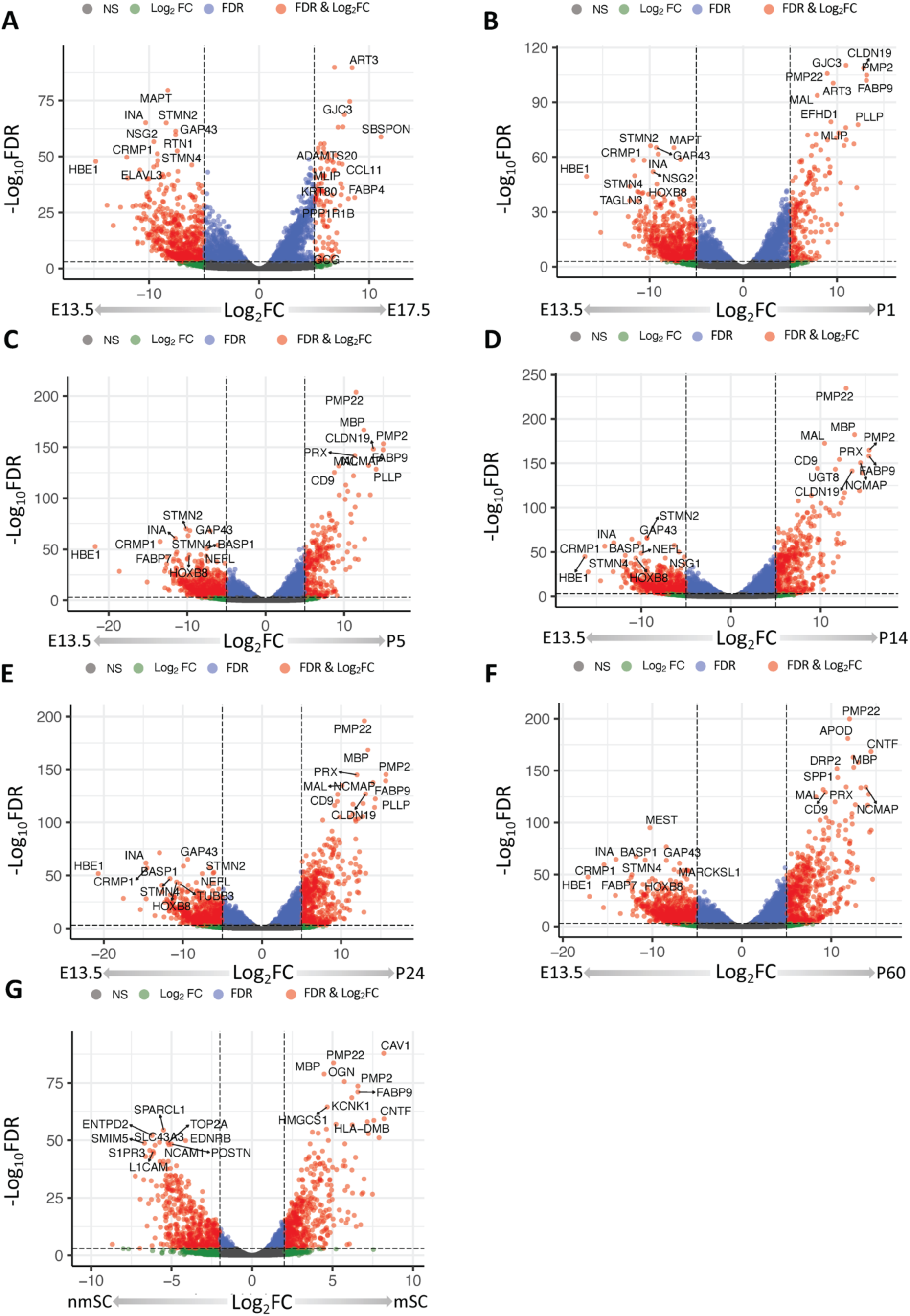
Differentially expression analysis of the SNAT RNA-seq dataset (GSE137868). Volcano plots depicting differentially expressed genes when comparing the E17.5 **(A)**, P1 **(B)**, P5 **(C)**, P14 **(D)**, P24 **(E)**, and P60 **(F)** stages to E13.5. Volcano plot depicting differentially expressed genes when comparing myelinating vs. non-myelinating Schwann cells **(G)**.

### Differential gene expression and enrichment in myelinating versus non-myelinating murine Schwann cells

We next sought to investigate the gene expression profiles of myelinating (mSCs) and non-myelinating (nmSCs) in the Sciatic Nerve Atlas. mSCs had 478 up-regulated DEGs and 471 down-regulated DEGs in comparison to nmSCs (**Fig. 4G, Table S5**). These up-regulated DEGs were enriched for structural constituents of myelin sheath (GO:0019911, *P=0.039*), myelination (GO:0042552, *P<0.001*), and sterol biosynthetic processes (GO:0016126, *P<0.001*). Down-regulated DEGs were highly enriched for mitotic cell cycle (GO:0000278, *P<0.001*), cell division (GO:0051301, *P<0.001),* and positive regulation of cell cycle processes (GO:0090068, *P<0.001*).

The myelinating and non-myelinating Schwann cell subsets remained largely distinct even upon recomputing the UMAP dimensionality reduction (**Fig. 5A**)[29]. Two distinct trajectories emerged, both beginning from proliferating Schwann cells (prol. SC) and terminating at either mSC cluster 3 or nm(R)SC (**Fig. 5A**)[29]. We observed 261 up-regulated and 116 down-regulated DEGs in myelinating Schwann cells (mSC) as compared to immature Schwann cells (iSCs) (**Table S6**). As expected, the up-regulated DEGs in mSCs were enriched for terms related to regulation of nervous system development (GO:0051960, *P<0.001*) and myelination (GO:0042552, *P<0.001*). However, the most significantly enriched term was sterol biosynthetic process (GO:0016126, *P<0.001*). The robust upregulation of cell cycle inhibitors like *CDKN1A* (**Fig. 5B**) in conjunction with the downregulation of cyclins such as *CDK6* (**Fig. 5B**) further suggest a state of proliferative arrest within this cluster. Using as few as three transcriptomic markers, we could distinguish the stages of Schwann cell differentiation and the bifurcation of mSCs and nmSCs (**Fig. 5C & D, Table S7**). Again, we identified ECM-related markers in the different differentiation stage iSC (*POSTN*), tSC (*ID3*), and mSC (*SPP1*) clusters (**Table S7**).

**Fig. 5.**
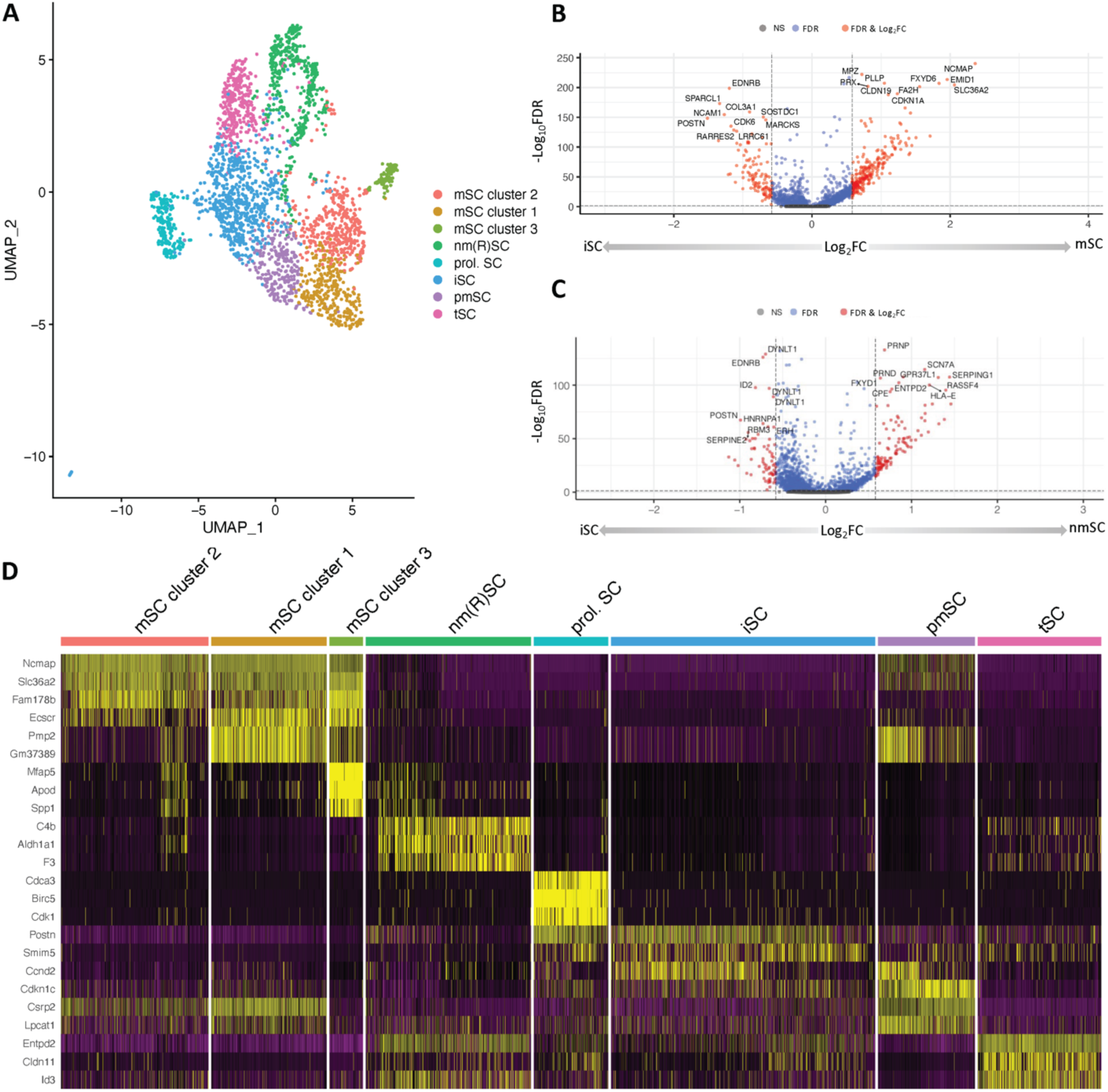
SNAT scRNA-seq analysis. **A)** Scatterplot of 8 clusters identified from the SNAT study representing different stages of Schwann cell development along both myelinating and non-myelinating trajectories. **B)** Volcano plot depicting differentially expressed genes comparing mSC vs iSC. **C)** Volcano plot depicting differentially expressed genes between nmSC versus iSC. **D)** Heatmap of cluster specific markers (top 3).

### Differentiating Schwann cells have conserved pathways related to ECM

As anticipated, the transcriptional signatures of Schwann cell differentiation were not identically conserved between hiPSCs and mice, likely due to both species and model specific differences. Without normalization, there was no significant correlation between the developmental stages within the SNAT dataset and the iPSC differentiation timepoints (**Fig. S22**). However, after accounting for differences in the row and column DEG counts, a moderate association emerged (PCC=0.44, *P=0.007*, **Fig 6A**). The one-to-one correspondences in DEGs between timepoints (E17.5-Day 3, P1-Day 6, P5-Day 12, P14-Day 18, P24-Day 24, and P60-Day 31) were also moderately associated (*P=0.084*, **Fig. 6B**).

**Fig. 6.**
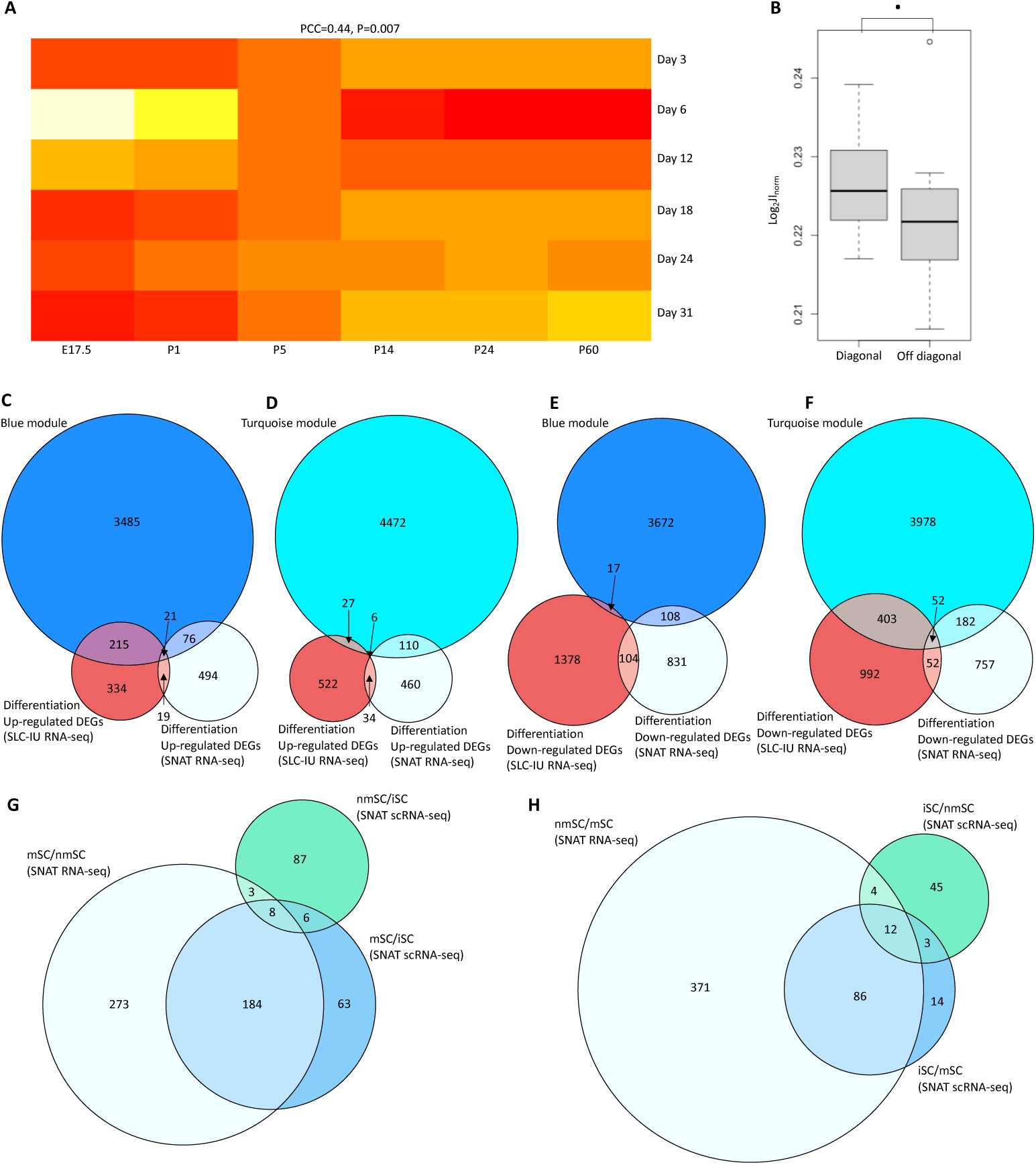
Comparison of DEGs identified between multiple datasets. **A)** Normalized JI matrix of averaged SC-IU DEGs (|log2FC| > 0.58 and FDR < 0.05) throughout SC differentiation and SNAT DEGs (|log2FC| > 0.58 and FDR < 0.05) through various embryonic and postnatal developmental states with comparison of the JI matrix diagonal to the non-diagonal **(B)**, **C)** Venn diagram comparing genes comprising the blue module, SC-IU up-regulated DEGs during SC differentiation, and SNAT up-regulated DEGs throughout development, **D)** Venn diagram comparing genes comprising the turquoise module, SC-IU up-regulated DEGs during SC differentiation, and SNAT up-regulated DEGs throughout development, **E)** Venn diagram comparing genes comprising the blue module, SC-IU down-regulated DEGs during SC differentiation, and SNAT down-regulated DEGs throughout development, **F)** Venn diagram comparing genes comprising the turquoise module, SC-IU down-regulated DEGs during SC differentiation, and SNAT down-regulated DEGs throughout development, **G)** Comparison of genes up-regulated in mSC vs. iSC and nmSC vs. iSC from SNAT scRNA-seq and DEGs upregulated in mSC vs. nmSC from SNAT RNA-seq, **H)** Comparison of genes down-regulated in mSCs vs. and nmSC vs. iSC from SNAT scRNA-seq and DEGs down-regulated in mSC vs. nmSC from SNAT RNA-seq.

We focused on the blue module and turquoise modules and compared them to the up- and down-regulated DEGs from the SC-IU and SNAT datasets. The blue module contained *NF2* and the eigengene trended upward throughout the differentiation process, whereas the turquoise module trended downward. We found that the up-regulated DEGs from both SC-IU and SNAT had high overlap with the blue module, but low overlap with one another (**Fig. 6C**). In contrast the SC-IU up-regulated DEGs had lower overlap with the turquoise module, while SNAT up-regulated DEGs had a similar number of DEGs in the turquoise module (**Fig. 6D**). Both the SC-IU and SNAT down-regulated DEGs had lower overlap with the blue module (**Fig. 6E**) than with the turquoise module (**Fig. 6F**). Notably 21 genes were up-regulated in both datasets and contained within the blue module (**Table S2**). These genes represented specific molecular functions including extracellular matrix structural constituent (GO:0005201, *P=0.005*) and S100 protein binding (GO:0044548, *P=0.008*) as well as biological processes including caveola assembly (GO:0070836, *P=0.006*) and regulation of transforming growth factor beta receptor signaling pathway (GO:0017015, *P=0.006*).

In addition to these unique conserved processes during murine Schwann cell development and hiPSC to Schwann cell differentiation, mSCs had distinct molecular profiles in comparison to nmSCs that were recapitulated in both RNA-seq and scRNA-seq data modalities. Notably, mSC up-regulated DEGs from SNAT RNA-seq had high overlap with mSC markers from SNAT scRNA-seq (**Fig. 6G**). Similarly, nmSC up-regulated DEGs from SNAT RNA-seq exhibited a high degree of overlap with nmSC markers from SNAT scRNA-seq (**Fig. 6H**).

## DISCUSSION

We have developed a state-of-the-art platform to study Schwann cell development using hiPSCs. We demonstrate that hiPSCs successfully differentiated into SCs based on marker gene expression and morphology. Furthermore, we identify similar transcriptomic profiles in our hiPSC to SC differentiation model as observed in other external Schwann cell differentiation studies [28, 29]. Notably, we observed correlation of DEGs and gene expression signatures including those relating to ECM and TGF-β signaling between hiPSC derived SCs and those arising in situ from the developing mouse sciatic nerve [29]. Intriguingly, TGF-β1 signaling has been shown to coordinate the production of abundant ECM proteins by *Nf1* deficient Schwann cells, contributing to plexiform neurofibroma pathogenesis in neurofibromatosis type 1 (NF1) [39]. However, the role of ECM and TGF-β signaling in schwannoma development remains largely understudied.

Key differences between our hiPSC model and mouse Schwann cell development also emerged, which may be attributable to both species-specific differences as well as the model system employed (*in vitro* versus *in vivo*). Transcriptional discordance between SCs derived from hAMSCs versus hiPSCs may in part be attributable to unique cells of origin and the experimental differences related the reprograming process. While hiPSCs have been utilized successfully to overcome limitation of mouse models to enhance our understanding of psychiatric disorders [40] and Alzheimer’s disease [21], further investigation is needed to validate the fidelity of this platform in recapitulate the pathogenesis of *NF2*-SWN and other Schwann cell associated disorders.

Notwitstanding these disparities, our analysis did reveal distinctive gene signatures associated with Schwann cell development that were conserved across species and model systems. We identified gene co-expression modules related to differentiation, particularly the ‘blue module’ that contained the *NF2* tumor suppressor gene. This module demonstrated functional enrichment for multiple terms related to ECM and actin filaments, which is consistent with the established relationship between Merlin and F-actin dependent cytoskeletal defects in *NF2*-associated schwannoma [41]. The emergence of ECM-related genes was a consistent finding across datasets, analytical techniques, and experimental methodologies employed in the study of differentiating Schwann cells. Notably, two vesicle-associated genes within the ‘blue module’, *CAV1* and *CAV2*, were found to be upregulated in both hiPSC and murine derived SCs and recently *NF2* was shown to regulate vesicle trafficking in schwannoma [42]. Additionally, collagens *COL1A1* and *COL3A1*, were also overrepresented in our analysis and recently shown to be associated with recurrence of vestibular schwannoma after radiation therapy [43]. Other ECM-related genes identified, albeit with lesser substantiation in schwannoma included *ITGA1*, *POSTN*, *SPP1*, *ID3*, and *ADAMTS6*. One of the most promising candidates was lysyl oxidase, i.e., *LOX*, which recurred in multiple analyses. LOX is an essential protein in the ECM that mediates conversion of collagen precursors into reactive forms capable of cross-linking and stabilizing ECM proteins to modulate cell adhesion and motility [44].

Cell cycle inhibitors, such as *CDKN2B*, were also present in the ‘blue module’ and conserved in cross species analysis. *CDKN2B* is a key component of the *INK4/ARF* tumor suppressor locus and can induce proliferation when deleted or mutated in a multitude of cancer types [45, 46]. Furthermore, we identified hiPSC-specific genes of interest that were highly up-regulated during SC differentiation and differentially expressed at all timepoints including *ANXA1,* which has been implicated in promoting Schwann cell proliferation and migration in response to nerve injury [47], yet its relationship to Merlin in Schwann cell biology remains unknown.

In summary, we developed a novel hiPSC Schwann cell differentiation model and performed integrated analyses across five datasets from three independent studies, encompassing both scRNA-seq and RNA-seq. Our findings indicate that our hiPSC model robustly recapitulates Schwann cell ontologies and identifies critical genes implicated in Schwann cell differentiation warranting further investigation. Additionally, our data revealed a pronounced association between SC development and ECM across multiple datasets, that remains largely uncharacterized with respect to Merlin biology and schwannoma development. Future studies leveraging this hiPSC platform to conduct CRISPR-Cas9 editing of *NF2* and other associated genes relevant to SC differentiation/function, generate hiPSC organoid co-cultures, and perform bulk and single cell mulitomic analysis hold promise to reveal new insights into Schwann cell biology, the Schwann cell-ECM relationship, and schwannomas pathogenesis in *NF2-*SWN.

## ACKNOWLEDGEMENTS

We would like to acknowledge the help and support of the Indiana University School of Medicine Center of Computational Biology and Bioinformatics, the Indiana University Information Technology Services for their high-performance computing resources, and the Indiana Biosciences Research Institute for their lab space and technical support.

## AUTHOR CONTRIBUTIONS

S.D.R, A.S.Q, and T.J. designed the experiments and evaluated the data. W.C, A.H, A.S.Q, and B.E.H. performed the experiments. O.L, S.L, and T.J analyzed all data and prepared the graph. T.J, S.L, A.S.Q, and S.D.R wrote the initial draft of the manuscript. S.P.A and D.W.C. gave advice during the project achievement. All co-authors provided feedback and revisions.

## Abbreviations

hiPSC: Human induced pluripotent stem cells
SC: Schwann cells
SPC: Schwann cell precursors
iSC: Immature SCs
mSC: Myelinating SCs
tSC: Transition SCs
pmSC: Pro-myelinating SCs
nmSC: Not-myelinating SCs
nm(R)SC: Non-myelinating (Remak) SCs

