## Supplementary Material for "A novel induced pluripotent stem cell model of Schwann cell differentiation reveals *NF2*- related gene regulatory networks of the extracellular matrix"

### TABLES

**Table S1** All significant marker DEGs for the clusters identified from the SC-SUMC dataset

| Gene | Cluster | Proportion cluster > 0 | Proportion other clusters >0 | $Log_2FC$ | P | FDR |
|---|---|---|---|---|---|---|
| *FTL* | 2 | 0.99 | 0.93 | 0.68 | 1.2E-58 | 1.4E-54 |
| *FTH1* | 2 | 1.00 | 0.98 | 0.60 | 7.2E-42 | 8.3E-38 |
| *CXCL1* | 3 | 0.96 | 0.79 | 1.15 | 1.5E-144 | 1.7E-140 |
| *TIMP1* | 3 | 0.89 | 0.60 | 1.34 | 8E-131 | 9.2E-127 |
| *TFPI2* | 3 | 0.79 | 0.47 | 1.12 | 2.2E-110 | 2.5E-106 |
| *CXCL8* | 3 | 0.87 | 0.60 | 1.02 | 7E-103 | 8E-99 |
| *B2M* | 3 | 0.64 | 0.34 | 0.68 | 2.5E-68 | 2.8E-64 |
| *SERPINE2* | 3 | 0.43 | 0.20 | 0.62 | 5.5E-45 | 6.3E-41 |
| *TAGLN* | 4 | 0.90 | 0.28 | 1.43 | 4.2E-242 | 4.8E-238 |
| *ACTA2* | 4 | 0.47 | 0.09 | 0.70 | 1.5E-133 | 1.7E-129 |
| *LGALS1* | 4 | 0.94 | 0.55 | 0.93 | 2.6E-125 | 3E-121 |
| *TPM1* | 4 | 0.80 | 0.36 | 1.02 | 4.9E-122 | 5.7E-118 |
| *TPM2* | 4 | 0.92 | 0.60 | 1.09 | 7.9E-121 | 9.1E-117 |
| *MYL9* | 4 | 0.57 | 0.18 | 0.59 | 1.4E-95 | 1.6E-91 |
| *TMSB10* | 4 | 0.99 | 0.89 | 0.68 | 7.1E-82 | 8.1E-78 |
| *ACTB* | 4 | 0.86 | 0.56 | 0.79 | 1.5E-77 | 1.8E-73 |
| *MYL6* | 4 | 0.90 | 0.64 | 0.62 | 4E-66 | 4.6E-62 |
| *RAB13* | 6 | 0.87 | 0.28 | 1.88 | 7.6E-144 | 8.8E-140 |
| *ANP32B* | 6 | 0.51 | 0.10 | 0.72 | 2E-96 | 2.3E-92 |
| *CCND1* | 6 | 0.41 | 0.09 | 0.72 | 1.9E-66 | 2.2E-62 |
| *IGFBP5* | 6 | 0.70 | 0.34 | 1.30 | 8E-50 | 9.2E-46 |
| *GAPDH* | 6 | 0.94 | 0.88 | 0.59 | 3.1E-17 | 3.5E-13 |
| *TPM2* | 6 | 0.76 | 0.62 | 0.68 | 4E-16 | 4.6E-12 |
| *LGALS1* | 6 | 0.69 | 0.58 | 0.59 | 1.6E-10 | 1.8E-06 |
| *ACTB* | 6 | 0.70 | 0.58 | 0.59 | 4.6E-10 | 5.3E-06 |
| *TAGLN* | 6 | 0.48 | 0.32 | 0.75 | 5.4E-10 | 6.2E-06 |
| *MALAT1* | 7 | 0.82 | 0.51 | 0.68 | 4.3E-40 | 4.9E-36 |
| *TUBA1B* | 8 | 0.68 | 0.18 | 0.95 | 1.5E-88 | 1.7E-84 |

| | | | | | | |
|---|---|---|---|---|---|---|
| *H2AFZ* | 8 | 0.73 | 0.27 | 0.87 | 2.3E-70 | 2.7E-66 |
| *UBE2S* | 8 | 0.50 | 0.12 | 0.66 | 1.2E-67 | 1.4E-63 |
| *LGALS1* | 8 | 0.88 | 0.58 | 0.64 | 2.4E-31 | 2.8E-27 |
| *VIM* | 8 | 0.79 | 0.52 | 0.74 | 2.3E-27 | 2.7E-23 |
| *ACTB* | 8 | 0.79 | 0.58 | 0.62 | 1.2E-19 | 1.4E-15 |
| *PPIB* | 10 | 0.42 | 0.16 | 0.59 | 3.8E-19 | 4.3E-15 |
| *IGFBP7* | 10 | 0.44 | 0.18 | 0.83 | 1.7E-17 | 2E-13 |
| *CD63* | 10 | 0.59 | 0.32 | 0.73 | 3.8E-16 | 4.4E-12 |
| *CXCL8* | 10 | 0.89 | 0.63 | 1.36 | 2.4E-14 | 2.7E-10 |
| *B2M* | 10 | 0.61 | 0.37 | 0.97 | 2.3E-13 | 2.6E-09 |
| *TIMP1* | 10 | 0.84 | 0.63 | 0.82 | 1.3E-12 | 1.5E-08 |
| *TFPI2* | 10 | 0.72 | 0.50 | 0.72 | 3.2E-12 | 3.6E-08 |
| *CXCL5* | 10 | 0.41 | 0.19 | 0.78 | 5.4E-12 | 6.2E-08 |
| *CXCL3* | 10 | 0.45 | 0.24 | 0.76 | 9.4E-10 | 1.1E-05 |
| *NEAT1* | 12 | 0.51 | 0.16 | 0.72 | 1.3E-25 | 1.5E-21 |
| *ITGB1* | 12 | 0.48 | 0.15 | 0.70 | 1.7E-22 | 1.9E-18 |
| *PAPPA* | 12 | 0.36 | 0.11 | 0.75 | 5.4E-18 | 6.2E-14 |
| *IGFBP5* | 12 | 0.65 | 0.35 | 0.70 | 2.6E-14 | 3E-10 |
| *MALAT1* | 12 | 0.70 | 0.52 | 0.93 | 1.1E-09 | 1.3E-05 |
| *KCNQ1OT1* | 14 | 0.35 | 0.01 | 0.74 | 7.4E-186 | 8.5E-182 |
| *N4BP2L2* | 14 | 0.42 | 0.02 | 0.68 | 6.7E-125 | 7.7E-121 |
| *ADAMTS6* | 14 | 0.32 | 0.01 | 0.72 | 3.5E-123 | 4.1E-119 |
| *PNISR* | 14 | 0.41 | 0.02 | 0.74 | 3.6E-116 | 4.2E-112 |
| *WSB1* | 14 | 0.42 | 0.02 | 0.87 | 1.2E-112 | 1.4E-108 |
| *MEG3* | 14 | 0.59 | 0.08 | 1.51 | 1E-72 | 1.2E-68 |
| *SLC25A37* | 14 | 0.36 | 0.03 | 0.77 | 6.8E-64 | 7.8E-60 |
| *NEAT1* | 14 | 0.73 | 0.15 | 4.25 | 4.5E-63 | 5.2E-59 |
| *ITGA1* | 14 | 0.39 | 0.04 | 0.93 | 5E-60 | 5.7E-56 |
| *DDX17* | 14 | 0.37 | 0.04 | 0.65 | 2.1E-54 | 2.4E-50 |
| *VMP1* | 14 | 0.48 | 0.08 | 1.60 | 7.3E-50 | 8.3E-46 |
| *MALAT1* | 14 | 0.91 | 0.52 | 5.12 | 7E-39 | 8E-35 |
| *COL1A1* | 14 | 0.58 | 0.14 | 0.74 | 4.6E-34 | 5.2E-30 |
| *CALD1* | 14 | 0.55 | 0.23 | 1.25 | 1.9E-18 | 2.2E-14 |

| | | | | | | |
|---|---|---|---|---|---|---|
| *TPM1* | 14 | 0.67 | 0.40 | 0.62 | 5.4E-09 | 6.2E-05 |
| *ACHE* | 15 | 0.14 | 0.00 | 0.90 | 3.1E-101 | 3.5E-97 |
| *GLCCI1* | 15 | 0.14 | 0.00 | 0.90 | 2E-39 | 2.3E-35 |
| *SMARCD3* | 15 | 0.23 | 0.01 | 0.82 | 5.8E-36 | 6.7E-32 |
| *TSC22D1* | 15 | 0.50 | 0.04 | 1.69 | 1.9E-26 | 2.1E-22 |
| *RFC2* | 15 | 0.18 | 0.01 | 0.62 | 6E-26 | 6.9E-22 |
| *AGAP3* | 15 | 0.23 | 0.02 | 0.61 | 6.5E-15 | 7.4E-11 |
| *BET1* | 15 | 0.36 | 0.04 | 0.81 | 9.8E-15 | 1.1E-10 |
| *TRIP6* | 15 | 0.27 | 0.02 | 0.60 | 1.4E-14 | 1.6E-10 |
| *ARL4A* | 15 | 0.18 | 0.01 | 0.61 | 1.2E-12 | 1.4E-08 |
| *DNAJC2* | 15 | 0.18 | 0.01 | 0.91 | 2.2E-12 | 2.5E-08 |
| *GNAI1* | 15 | 0.27 | 0.04 | 0.81 | 6.7E-09 | 7.6E-05 |
| *TSC22D4* | 15 | 0.18 | 0.02 | 0.84 | 1.6E-08 | 1.8E-04 |
| *AKAP9* | 15 | 0.14 | 0.01 | 1.08 | 1.6E-08 | 1.8E-04 |
| *SNHG15* | 15 | 0.14 | 0.01 | 1.11 | 2.8E-08 | 3.3E-04 |
| *UBE2H* | 15 | 0.23 | 0.03 | 0.78 | 4.2E-07 | 4.8E-03 |
| *WBSCR22* | 15 | 0.18 | 0.02 | 0.83 | 6.2E-07 | 7.1E-03 |
| *RHEB* | 15 | 0.46 | 0.12 | 1.34 | 8E-07 | 9.2E-03 |
| *ZYX* | 15 | 0.23 | 0.04 | 0.91 | 2.5E-06 | 2.8E-02 |
| *CAV1* | 15 | 0.18 | 0.03 | 0.99 | 3.4E-06 | 3.9E-02 |
| *SBDS* | 15 | 0.27 | 0.06 | 0.78 | 4.3E-06 | 4.9E-02 |

**Table S2** DGE analysis of Schwann cell differentiation in the SNAT RNA-seq dataset. The top 5 up- and down-regulated DEGs for each of the 6 subsequent stages (k) compared to E13.5. Gene names are capitalized because they have been converted to their human homologs (See **METHODS**).

| Stage k vs. Stage E13.5 | Gene | $Log_2FC$ | P | FDR |
|---|---|---|---|---|
| E17.5 Up | *MAL* | 6.84 | 6.86E-95 | 1.42E-90 |
| E17.5 Up | *ART3* | 8.44 | 2.00E-94 | 2.08E-90 |
| E17.5 Up | *GJC3* | 8.22 | 5.23E-79 | 2.71E-75 |

| | | | | |
|---|---|---|---|---|
| E17.5 Up | *LAMA2* | 7.16 | 3.07E-67 | 7.08E-64 |
| E17.5 Up | *SBSPON* | 11.07 | 9.40E-63 | 1.63E-59 |
| E17.5 Down | *MAPT* | -8.30 | 4.04E-84 | 2.79E-80 |
| E17.5 Down | *INA* | -10.32 | 2.09E-69 | 7.24E-66 |
| E17.5 Down | *STMN2* | -8.45 | 2.91E-69 | 8.62E-66 |
| E17.5 Down | *GAP43* | -7.60 | 2.09E-65 | 4.34E-62 |
| E17.5 Down | *RTN1* | -7.61 | 1.05E-63 | 1.98E-60 |
| P1 Up | *GJC3* | 10.96 | 2.48E-115 | 5.15E-111 |
| P1 Up | *CLDN19* | 12.85 | 1.49E-113 | 1.55E-109 |
| P1 Up | *PMP22* | 8.96 | 2.36E-110 | 1.63E-106 |
| P1 Up | *PMP2* | 13.17 | 2.42E-109 | 1.26E-105 |
| P1 Up | *FABP9* | 13.15 | 2.15E-106 | 8.91E-103 |
| P1 Down | *STMN2* | -9.90 | 6.00E-70 | 6.23E-67 |
| P1 Down | *GAP43* | -9.27 | 6.48E-69 | 6.12E-66 |
| P1 Down | *MAPT* | -7.39 | 7.38E-69 | 6.66E-66 |
| P1 Down | *DPYSL3* | -6.70 | 3.11E-62 | 2.30E-59 |
| P1 Down | *CRMP1* | -11.78 | 8.43E-62 | 5.75E-59 |
| P5 Up | *PMP22* | 11.53 | 8.87E-209 | 1.84E-204 |
| P5 Up | *MBP* | 12.53 | 1.73E-171 | 1.80E-167 |
| P5 Up | *PMP2* | 15.02 | 6.84E-158 | 4.73E-154 |
| P5 Up | *CLDN19* | 13.74 | 1.44E-152 | 7.48E-149 |
| P5 Up | *FABP9* | 14.99 | 7.22E-152 | 3.00E-148 |
| P5 Down | *STMN2* | -10.12 | 1.96E-73 | 1.23E-70 |
| P5 Down | *GAP43* | -9.66 | 1.09E-71 | 6.44E-69 |
| P5 Down | *DPYSL3* | -7.07 | 2.56E-71 | 1.43E-68 |

| | | | | |
|---|---|---|---|---|
| P5 Down | *INA* | -11.51 | 4.25E-64 | 1.88E-61 |
| P5 Down | *CRMP1* | -13.45 | 7.73E-61 | 2.97E-58 |
| P14 Up | *PMP22* | 12.84 | 1.34E-239 | 2.77E-235 |
| P14 Up | *MBP* | 13.79 | 7.41E-187 | 7.69E-183 |
| P14 Up | *MAL* | 10.44 | 4.76E-177 | 3.29E-173 |
| P14 Up | *PMP2* | 15.41 | 2.11E-169 | 1.09E-165 |
| P14 Up | *FABP9* | 15.39 | 1.32E-162 | 5.49E-159 |
| P14 Down | *GAP43* | -9.38 | 2.39E-69 | 8.56E-67 |
| P14 Down | *STMN2* | -9.36 | 5.84E-69 | 2.04E-66 |
| P14 Down | *INA* | -13.07 | 1.19E-63 | 3.21E-61 |
| P14 Down | *BASP1* | -10.25 | 4.51E-63 | 1.14E-60 |
| P14 Down | *DPYSL3* | -6.31 | 1.22E-60 | 2.85E-58 |
| P24 Up | *PMP22* | 12.93 | 5.98E-201 | 1.24E-196 |
| P24 Up | *MBP* | 13.38 | 3.01E-173 | 3.12E-169 |
| P24 Up | *PMP2* | 15.66 | 7.27E-150 | 5.03E-146 |
| P24 Up | *PRX* | 12.00 | 2.38E-149 | 1.24E-145 |
| P24 Up | *FABP9* | 15.63 | 1.24E-143 | 5.16E-140 |
| P24 Down | *GAP43* | -9.38 | 1.71E-68 | 6.69E-66 |
| P24 Down | *INA* | -14.66 | 5.67E-65 | 1.93E-62 |
| P24 Down | *BASP1* | -9.92 | 4.82E-62 | 1.49E-59 |
| P24 Down | *MEST* | -6.56 | 7.22E-61 | 2.14E-58 |
| P24 Down | *CRMP1* | -14.53 | 3.92E-60 | 1.13E-57 |
| P60 Up | *PMP22* | 12.03 | 6.44E-205 | 1.34E-200 |
| P60 Up | *APOD* | 11.84 | 9.53E-186 | 9.89E-182 |
| P60 Up | *CNTF* | 14.44 | 9.13E-173 | 6.31E-169 |

| | | | | |
|---|---|---|---|---|
| P60 Up | *MBP* | 12.50 | 1.71E-157 | 5.93E-154 |
| P60 Up | *DRP2* | 10.64 | 3.58E-156 | 1.06E-152 |
| P60 Down | *MEST* | -10.27 | 1.21E-98 | 8.67E-96 |
| P60 Down | *INA* | -14.05 | 8.93E-68 | 2.52E-65 |
| P60 Down | *BASP1* | -10.80 | 4.56E-67 | 1.23E-64 |
| P60 Down | *GAP43* | -8.43 | 8.41E-67 | 2.21E-64 |
| P60 Down | *DPYSL3* | -6.96 | 6.41E-64 | 1.53E-61 |

**Table S3** Significant up and down regulated DEGs identified from SC-IU and SNAT datasets. SNAT gene names are capitalized because they have been converted to their human homologs (See **METHODS**).

| Gene set | Genes |
|---|---|
| Significantly up-regulated DEGs in all six comparisons for SC-IU dataset | *ACTA2, ANKRD1, ANXA1, CDH6, FILIP1L, GREM1, HAND1, LRRN3, LUM, MEIS2, NPPB, NPR3, NR2F1, NR2F2, SNAI2, TNFRSF19, WNT2B, ZEB2* |
| Significantly down-regulated DEGs at each timepoint comparison for SC-IU dataset | *AC003975.1, AC004543.1, AC007001.1, AC009446.1, AC022140.1, AC022140.2, AC073071.1, AC097520.1, AC104257.1, AC104758.3, AC108515.1, AC148477.4, AL117378.1, AL117378.2, AL157817.1, AL354821.1, AL354994.1, AL392023.1, AL591030.1, AP000943.2, B3GNT7, C10orf142, CR2, CTCFL, D21S2088E, DPEP3, EMX1, GFY, GLB1L3, GOLGA6A, GOLGA6C, GOLGA6D, GUCY2C, HCG24, HMX2, HRC, LCK, LINC00599, LINC00678, LINC01194, LNCPRESS2, MACC1-AS1, MAT1A, OLIG1, SLC35D3, TDGF1, TNFSF11, TRDN, UCMA, VWC2* |
| Significantly up-regulated DEGs at each timepoint comparison for SNAT dataset | *ADAMTS20, ART3, ASPA, BCAS1, C1S, CASP12, CCL11, CDH19, CLDN19, COL5A3, CRYAB, CTNNA3, CYP2J2, DEPDC7, DMD, EFHD1, EGR2, EMP2, ENTPD2, FABP4, GAL3ST1, GAS2L3, GATM, GJC3, IGFBP6, ITGB4, ITIH5, MAL, MBP, MLIP, NIPAL1, PLLP, PLP1, PLXDC1, PPP1R1B, PREX2, PRRG4, PRX, RARRES2, REM1, S100A3, S100A4, S100A6, SBSPON, SCN7A, THBS4, UGT8* |

| | |
|---|---|
| Significantly up-regulated DEGs in both SC-IU, SNAT, and contained in the blue module | *CCDC3, CDKN2B, COL8A1, CAV1, CAV2, KRT80, LOX, MFAP5, NNMT, NT5E, COL1A1, AHNAK, IGFBP7, LIMS2, MGLL, PALMD, SRGN, BHLHE40, FGF1, FNDC1, and FOXS1* |

**Table S4** Table of the modules that changed significantly in relation to developmental stage in the SNAT dataset evaluated by ANOVA. The number of genes in the module, ANOVA P, and most significant GO term are listed for each module.

| Module | Genes | P | Top GO term |
|---|---|---|---|
| Black | 305 | 3.39E-16 | Small GTPase binding |
| Blue | 2263 | 3.11E-11 | Positive regulation of DNA metabolic process |
| Brown | 2097 | 4.36E-6 | Regulation of neuron projection development |
| Green | 773 | 7.38E-5 | NADP metabolic process |
| Grey | 8 | 4.82E-8 | Proteasome-mediated ubiquitin-dependent protein catabolic process |
| Purple | 132 | 5.31E-7 | Purine-containing compound biosynthetic process |
| Tan | 92 | 3.49E-10 | Phagocytic cup |
| Turquoise | 4473 | 1.61E-10 | Regulation of protein stability |
| Yellow | 1246 | 2.10E-22 | Mitochondrial matrix |

**Table S5** DGE analysis results of the mSC and nmSC SNAT RNA-seq dataset including the top 10 up- and down-regulated DEGs in mSCs versus nmSCs from the SNAT RNA-seq dataset. Gene names are capitalized because they have been converted to their human homologs (See **METHODS**).

| Gene | $Log_2FC$ | P | FDR |
|---|---|---|---|
| *CAV1* | 8.18 | 1.14E-92 | 1.46E-88 |

| | | | |
|---|---|---|---|
| *PMP22* | 5.06 | 2.99E-88 | 1.92E-84 |
| *MBP* | 4.48 | 3.72E-83 | 1.59E-79 |
| *OGN* | 5.73 | 8.09E-80 | 2.59E-76 |
| *PMP2* | 6.57 | 8.89E-78 | 2.28E-74 |
| *FABP9* | 6.57 | 5.10E-75 | 1.09E-71 |
| *KCNK1* | 6.19 | 1.64E-72 | 3.01E-69 |
| *HMGCS1* | 4.67 | 1.72E-68 | 2.76E-65 |
| *CNTF* | 8.20 | 3.32E-63 | 4.73E-60 |
| *HLA-DMB* | 7.56 | 1.71E-62 | 2.19E-59 |
| *SPARCL1* | -5.50 | 4.07E-58 | 3.48E-55 |
| *ENTPD2* | -6.21 | 1.17E-55 | 7.91E-53 |
| *EDNRB* | -4.13 | 2.47E-53 | 1.44E-50 |
| *TOP2A* | -5.22 | 2.05E-52 | 1.10E-49 |
| *SMIM5* | -6.69 | 3.69E-52 | 1.89E-49 |
| *POSTN* | -5.01 | 9.13E-52 | 4.19E-49 |
| *NCAM1* | -5.16 | 1.82E-51 | 8.05E-49 |
| *SLC43A3* | -6.03 | 3.43E-51 | 1.47E-48 |
| *S1PR3* | -6.36 | 5.58E-49 | 2.31E-46 |
| *L1CAM* | -6.15 | 1.87E-48 | 7.27E-46 |

**Table S6** Top 10 up- and down-regulated DEGs comparing mSC versus iSC and nmSC versus iSC in the SNAT dataset. Gene names are capitalized because they have been converted to their human homologs (See **METHODS**).

| Cluster vs iSC | Gene | Proportion cluster > 0 | Proportion other clusters >0 | $Log_2FC$ | P | FDR |
|---|---|---|---|---|---|---|
| mSC Up | *NCMAP* | 0.96 | 0.18 | 2.36 | 3.91E-245 | 6.14E-241 |

| | | | | | | |
|---|---|---|---|---|---|---|
| mSC Up | *MPZ* | 1.00 | 0.99 | 0.72 | 6.62E-227 | 1.04E-222 |
| mSC Up | *EMID1* | 0.92 | 0.17 | 1.96 | 2.48E-218 | 3.90E-214 |
| mSC Up | *PLLP* | 1.00 | 0.65 | 1.05 | 3.71E-212 | 5.83E-208 |
| mSC Up | *FXYD6* | 0.93 | 0.22 | 1.84 | 7.08E-212 | 1.11E-207 |
| mSC Up | *SLC36A2* | 0.88 | 0.09 | 2.06 | 3.76E-209 | 5.90E-205 |
| mSC Up | *PRX* | 0.99 | 0.78 | 0.81 | 8.18E-207 | 1.28E-202 |
| mSC Up | *FA2H* | 0.96 | 0.27 | 1.56 | 2.50E-206 | 3.92E-202 |
| mSC Up | *CLDN19* | 0.99 | 0.48 | 1.24 | 1.86E-194 | 2.91E-190 |
| mSC Up | *CDKN1A* | 0.98 | 0.57 | 1.11 | 8.05E-193 | 1.26E-188 |
| mSC Down | *EDNRB* | 0.87 | 1.00 | -1.19 | 1.38E-203 | 2.16E-199 |
| mSC Down | *SPARCL1* | 0.59 | 0.96 | -1.34 | 5.24E-178 | 8.22E-174 |
| mSC Down | *COL3A1* | 0.92 | 1.00 | -0.90 | 8.02E-164 | 1.26E-159 |
| mSC Down | *NCAM1* | 0.51 | 0.95 | -1.27 | 1.79E-159 | 2.80E-155 |
| mSC Down | *SOSTDC1* | 0.94 | 0.98 | -0.70 | 2.53E-155 | 3.98E-151 |
| mSC Down | *POSTN* | 0.37 | 0.85 | -1.51 | 2.66E-153 | 4.17E-149 |
| mSC Down | *MARCKS* | 0.92 | 1.00 | -0.67 | 1.18E-150 | 1.85E-146 |
| mSC Down | *CDK6* | 0.60 | 0.91 | -1.17 | 4.10E-140 | 6.44E-136 |
| mSC Down | *RARRES2* | 0.64 | 0.89 | -1.13 | 1.11E-133 | 1.74E-129 |
| mSC Down | *LRRC61* | 0.55 | 0.88 | -1.09 | 1.38E-131 | 2.17E-127 |
| nmSC Up | *PRNP* | 0.98 | 0.81 | 0.69 | 9.16E-138 | 1.44E-133 |
| nmSC Up | *SCN7A* | 0.89 | 0.44 | 1.15 | 1.61E-119 | 2.52E-115 |
| nmSC Up | *PRND* | 0.90 | 0.61 | 0.90 | 1.64E-112 | 2.57E-108 |
| nmSC Up | *SERPING1* | 0.75 | 0.19 | 1.44 | 1.90E-112 | 2.99E-108 |
| nmSC Up | *GPR37L1* | 0.79 | 0.24 | 1.31 | 4.24E-112 | 6.66E-108 |
| nmSC Up | *FXYD1* | 0.98 | 0.84 | 0.64 | 1.41E-111 | 2.21E-107 |

| | | | | | | |
|---|---|---|---|---|---|---|
| nmSC Up | *ENTPD2* | 0.91 | 0.59 | 0.85 | 3.08E-107 | 4.84E-103 |
| nmSC Up | *HLA-E* | 0.83 | 0.32 | 1.21 | 5.46E-105 | 8.58E-101 |
| nmSC Up | *CPE* | 0.96 | 0.66 | 0.77 | 2.81E-101 | 4.41E-97 |
| nmSC Up | *RASSF4* | 0.67 | 0.13 | 1.40 | 2.84E-100 | 4.45E-96 |
| nmSC Down | *DYNLT1* | 0.83 | 0.98 | -0.70 | 7.57E-134 | 1.19E-129 |
| nmSC Down | *EDNRB* | 0.95 | 1.00 | -0.73 | 4.98E-131 | 7.82E-127 |
| nmSC Down | *ID2* | 0.89 | 0.98 | -0.82 | 1.31E-102 | 2.05E-98 |
| nmSC Down | *DYNLT1* | 0.81 | 0.96 | -0.66 | 5.92E-102 | 9.29E-98 |
| nmSC Down | *DYNLT1* | 0.85 | 0.97 | -0.61 | 5.11E-94 | 8.03E-90 |
| nmSC Down | *POSTN* | 0.60 | 0.85 | -0.99 | 2.58E-72 | 4.06E-68 |
| nmSC Down | *HNRNPA1* | 0.69 | 0.95 | -0.73 | 5.28E-69 | 8.29E-65 |
| nmSC Down | *RBM3* | 0.71 | 0.91 | -0.67 | 5.24E-66 | 8.23E-62 |
| nmSC Down | *ERH* | 0.78 | 0.95 | -0.60 | 1.05E-65 | 1.65E-61 |
| nmSC Down | *SERPINE2* | 0.78 | 0.88 | -0.90 | 9.23E-61 | 1.45E-56 |

**Table S7** Markers genes for SNAT scRNAseq dataset clusters.

| Gene | Cluster | Proportion cluster > 0 | Proportion other clusters >0 | $Log_2FC$ | P | FDR |
|---|---|---|---|---|---|---|
| *NCMAP* | mSC cluster 2 | 0.99 | 0.51 | 1.36 | 5.56E-157 | 8.74E-153 |
| *SLC36A2* | mSC cluster 2 | 0.92 | 0.38 | 1.26 | 1.29E-128 | 2.03E-124 |
| *FAM178B* | mSC cluster 2 | 0.79 | 0.28 | 1.31 | 4.81E-113 | 7.55E-109 |
| *ECSCR* | mSC cluster 1 | 0.85 | 0.19 | 1.45 | 1.77E-149 | 2.77E-145 |
| *PMP2* | mSC cluster 1 | 0.93 | 0.34 | 1.70 | 1.34E-143 | 2.10E-139 |
| *FABP9* | mSC cluster 1 | 0.92 | 0.33 | 1.69 | 1.64E-143 | 2.57E-139 |
| *MFAP5* | mSC cluster 3 | 1.00 | 0.10 | 2.48 | 2.20E-158 | 3.46E-154 |
| *APOD* | mSC cluster 3 | 0.99 | 0.12 | 2.29 | 2.43E-136 | 3.81E-132 |

| | | | | | | |
|---|---|---|---|---|---|---|
| *SPP1* | mSC cluster 3 | 0.89 | 0.10 | 2.26 | 3.41E-124 | 5.36E-120 |
| *C4A* | nm(R)SC | 0.69 | 0.09 | 1.70 | 3.53E-206 | 5.54E-202 |
| *ALDH1A1* | nm(R)SC | 0.65 | 0.08 | 1.70 | 2.20E-200 | 3.45E-196 |
| *F3* | nm(R)SC | 0.65 | 0.09 | 1.66 | 1.17E-182 | 1.83E-178 |
| *CDCA3* | prol. SC | 0.88 | 0.08 | 2.13 | 9.87E-245 | 1.55E-240 |
| *BIRC5* | prol. SC | 0.97 | 0.14 | 2.29 | 3.99E-223 | 6.27E-219 |
| *CDK1* | prol. SC | 0.87 | 0.16 | 2.17 | 3.47E-165 | 5.44E-161 |
| *POSTN* | iSC | 0.85 | 0.51 | 1.04 | 3.10E-116 | 4.87E-112 |
| *SMIM5* | iSC | 0.48 | 0.24 | 0.94 | 1.02E-47 | 1.60E-43 |
| *CCND2* | iSC | 0.58 | 0.36 | 0.89 | 7.06E-45 | 1.11E-40 |
| *CDKN1C* | pmSC | 0.96 | 0.51 | 1.46 | 7.81E-113 | 1.23E-108 |
| *CSRP2* | pmSC | 0.99 | 0.59 | 1.02 | 2.91E-77 | 4.58E-73 |
| *LPCAT1* | pmSC | 0.92 | 0.55 | 1.00 | 2.10E-74 | 3.30E-70 |
| *ENTPD2* | tSC | 0.97 | 0.53 | 1.19 | 5.28E-125 | 8.29E-121 |
| *CLDN11* | tSC | 0.66 | 0.14 | 1.14 | 8.61E-117 | 1.35E-112 |
| *ID3* | tSC | 0.77 | 0.31 | 1.20 | 2.69E-83 | 4.22E-79 |

### FIGURES

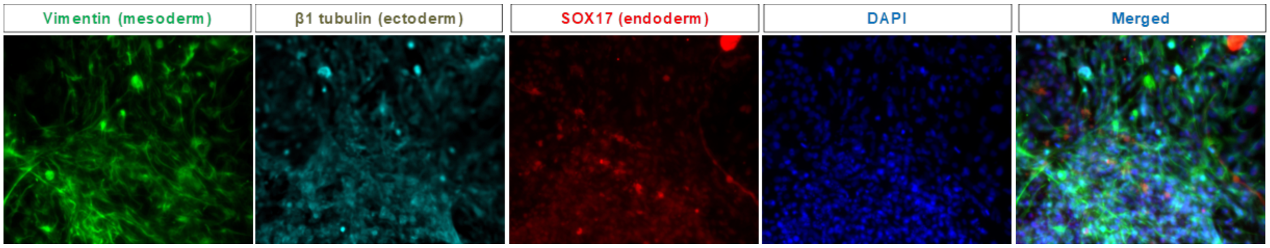


**Fig. S1** Characterization of embryoid body (EB) and three germ layer formation of iPSCs IBRI-101. J. Immunofluorescence microscopy of EBs merged with DAPI (in blue) showing expression of Vimentin (mesoderm), β1 tubulin (ectoderm) and SOX17 (endoderm).

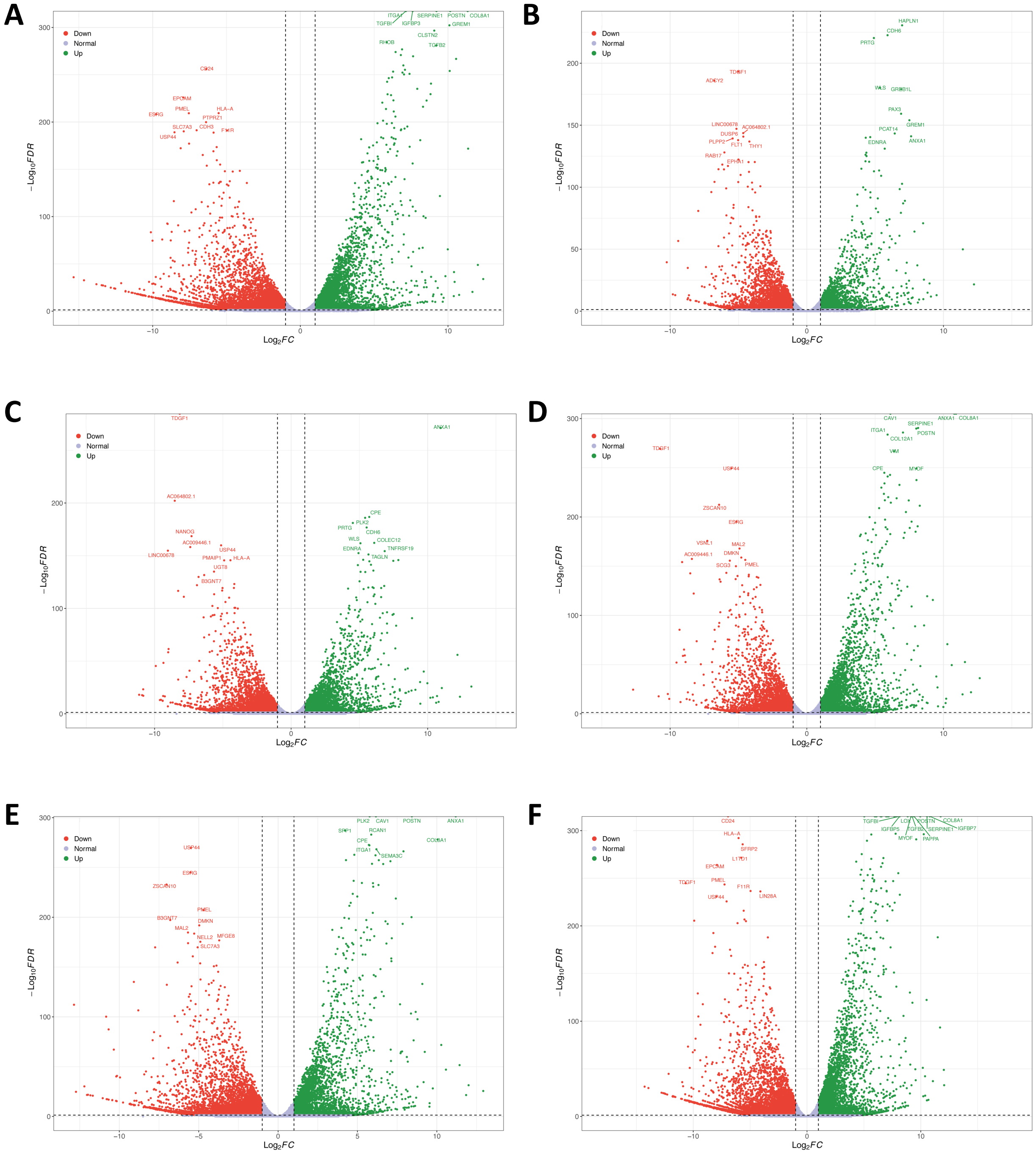


**Fig. S2 Volcano plots of differentially expressed genes during iPSC to SC differentiation from bulk RNAseq analysis of the SC-IU dataset.** Plots depict DEGs when comparing day 3 **(A)**, day 6 **(B)**, day 12 **(C)**, day 18 **(D)**, day 24 **(E)**, and day 31 **(F)** of SC differentiation to day 0 hiPSCs as the reference timepoint**.**

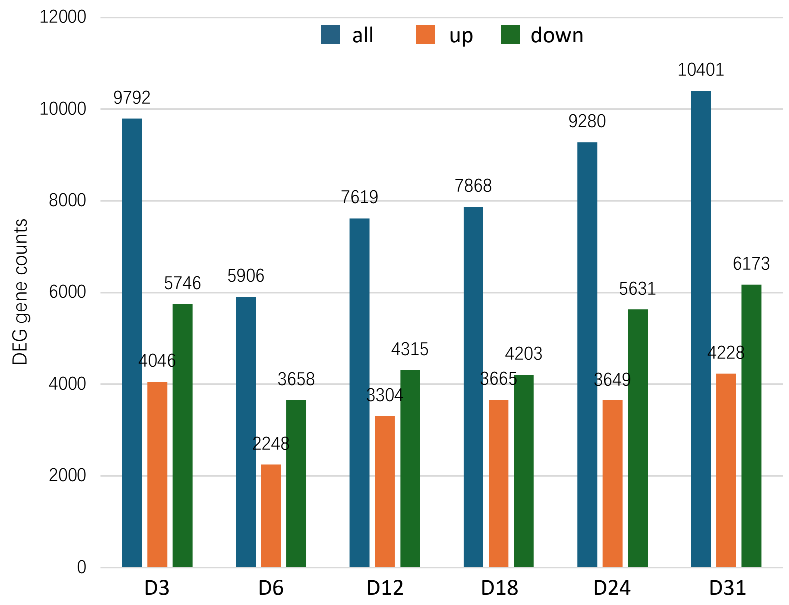


**Fig. S3** The number of DEGs for each comparison of SCs at various stages of differentiation (day 3, 6, 12, 18, 24, 31) compared to day 0 hiPSCs as a reference timepoint. Type denotes whether the DEG was up-regulated (up), down-regulated (down), or either (all).

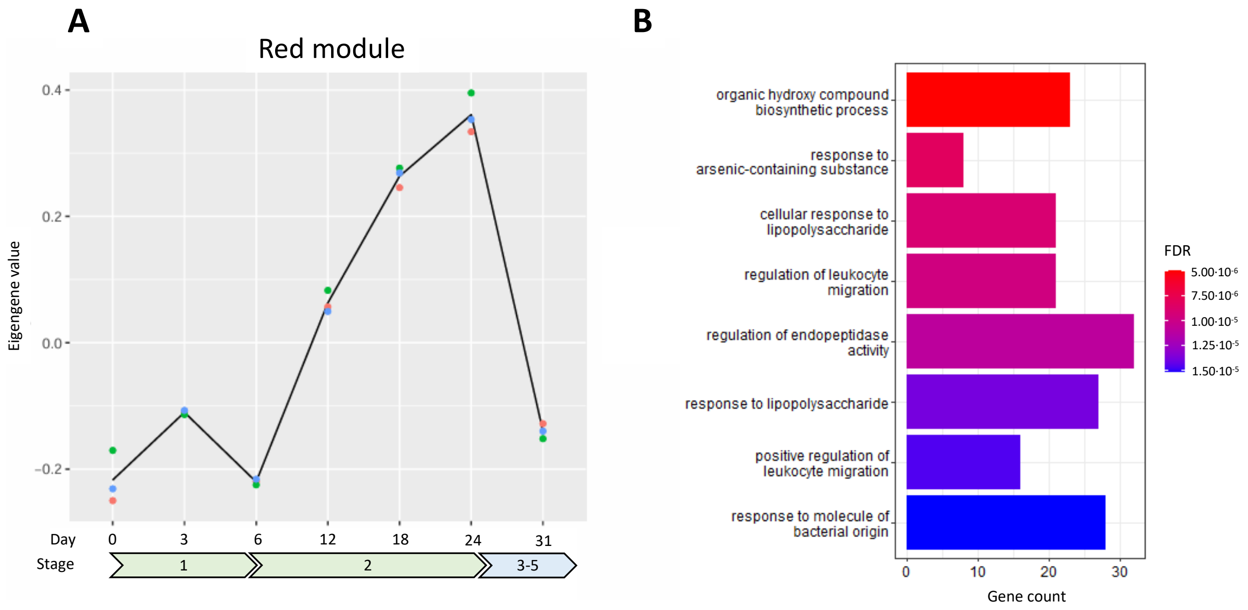


**Fig. S4** SC-IU red module results. **A)** Module eigengene are plotted over the course of hiPSC to SC differentiation. **B)** Functional enrichment analysis reveals significantly enriched ontology terms.

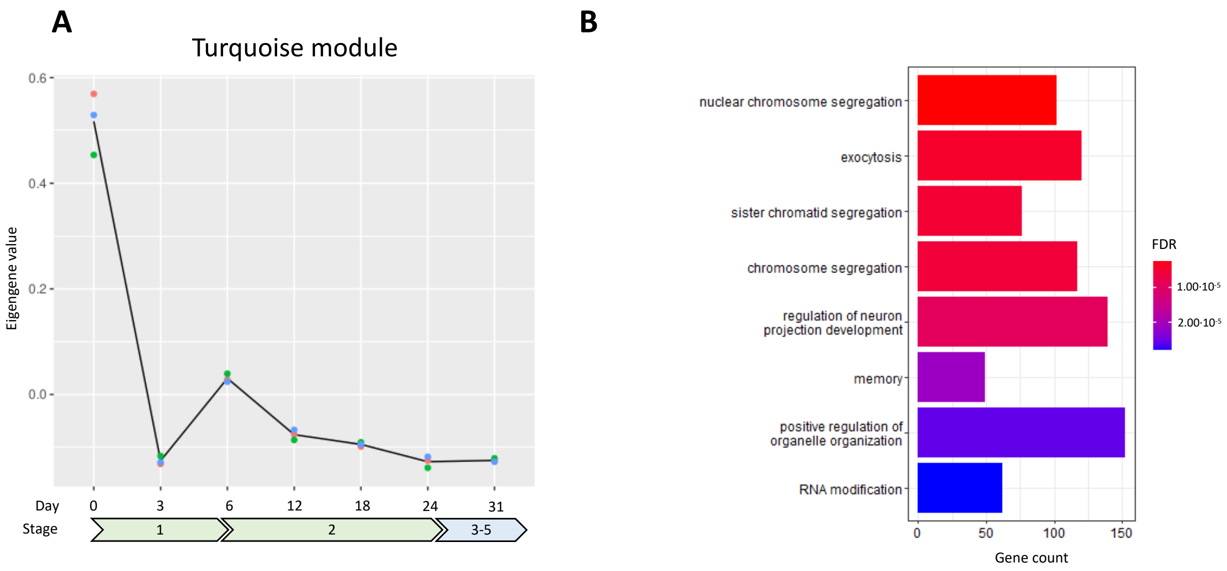


**Fig. S5** SC-IU turquoise module results. **A)** Module eigengene are plotted over the course of hiPSC to SC differentiation. **B)** Functional enrichment analysis reveals significantly enriched ontology terms.

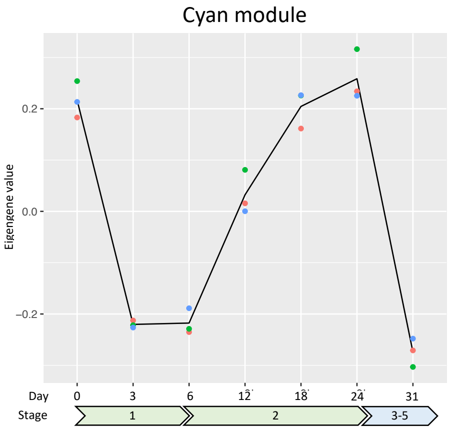


**Fig. S6** SC-IU cyan module results. Module eigengene are plotted over the course of hiPSC to SC differentiation. Functional enrichment analysis did not reveal significantly enriched ontology terms.

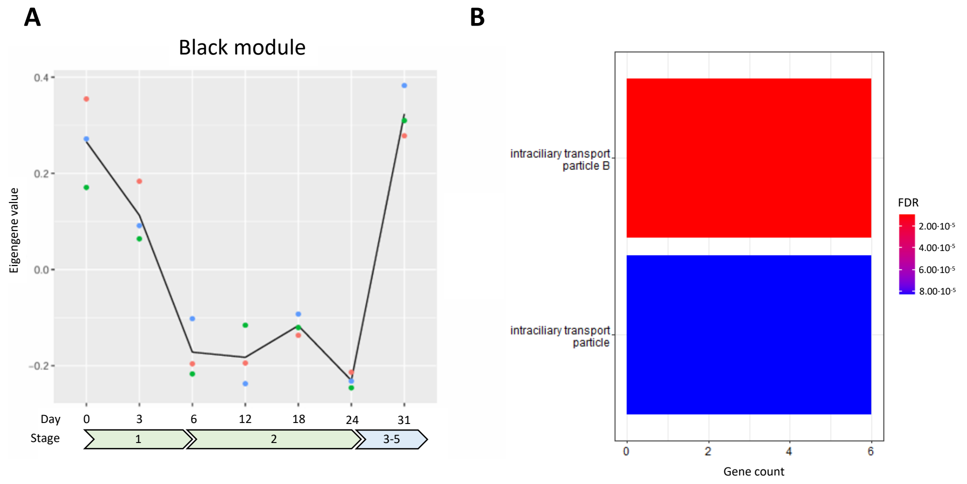


**Fig. S7** SC-IU black module results. **A)** Module eigengene are plotted over the course of hiPSC to SC differentiation. **B)** Functional enrichment analysis reveals significantly enriched ontology terms.

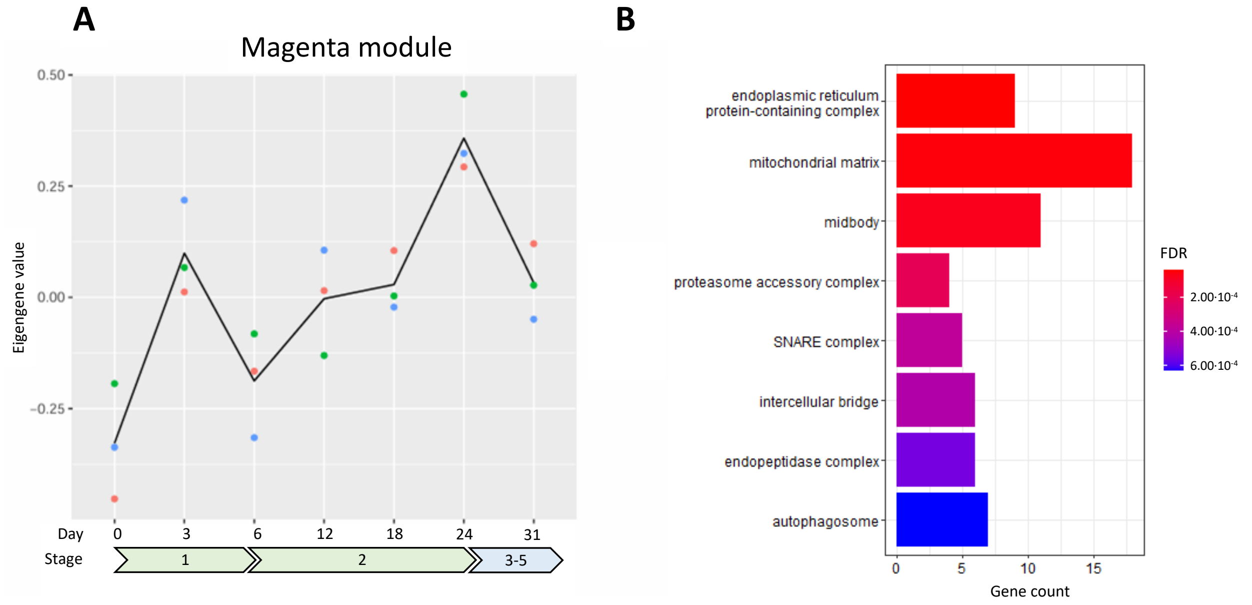


**Fig. S8** SC-IU magenta module results. **A)** Module eigengene are plotted over the course of hiPSC to SC differentiation. **B)** Functional enrichment analysis reveals significantly enriched ontology terms.

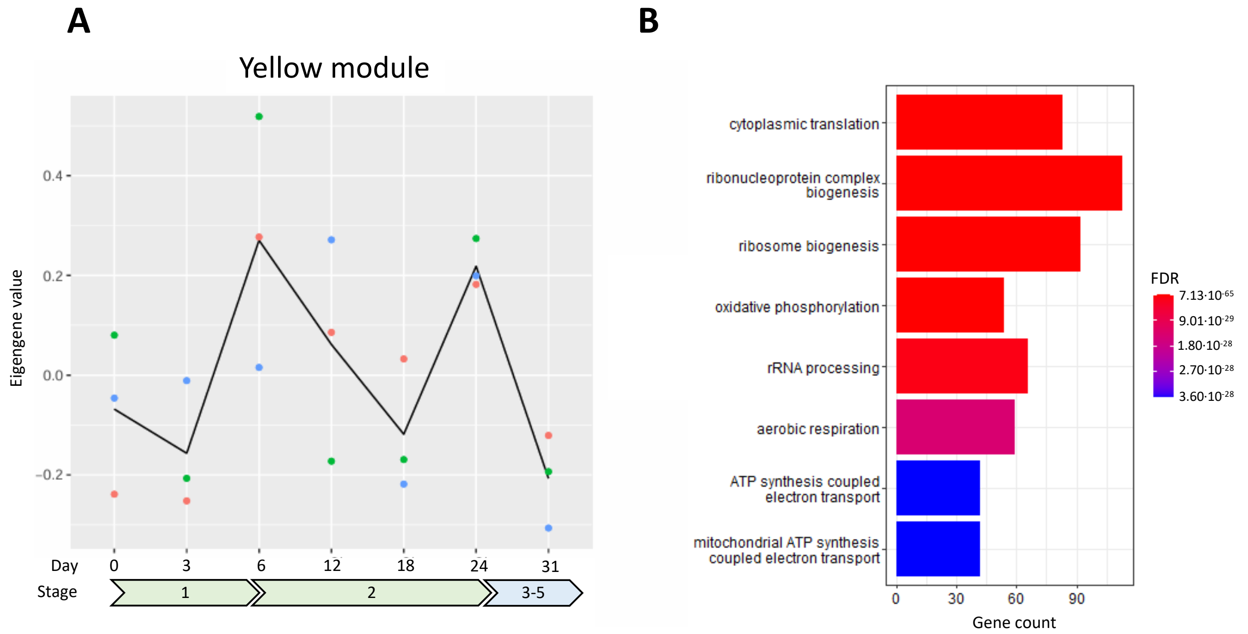


**Fig. S9** SC-IU **y**ellow module results. **A)** Module eigengene are plotted over the course of hiPSC to SC differentiation. **B)** Functional enrichment analysis reveals significantly enriched ontology terms.

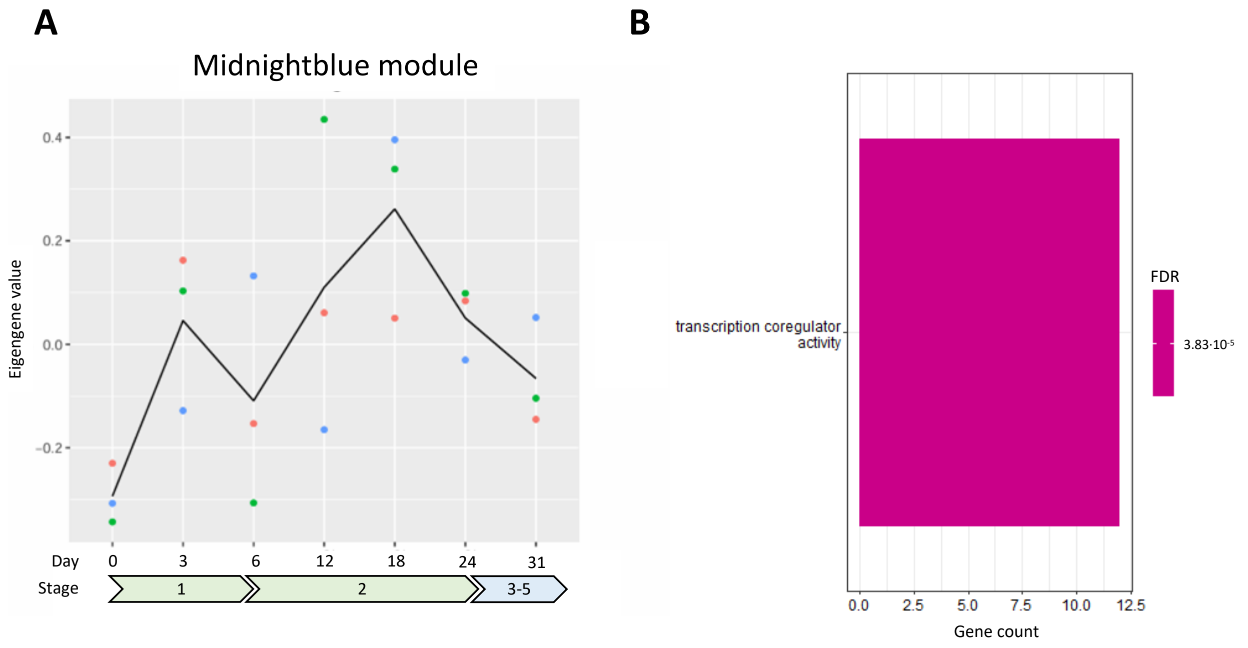


**Fig. S10** SC-IU midnightblue module results. **A)** Module eigengene are plotted over the course of hiPSC to SC differentiation. **B)** Functional enrichment analysis reveals significantly enriched ontology terms.

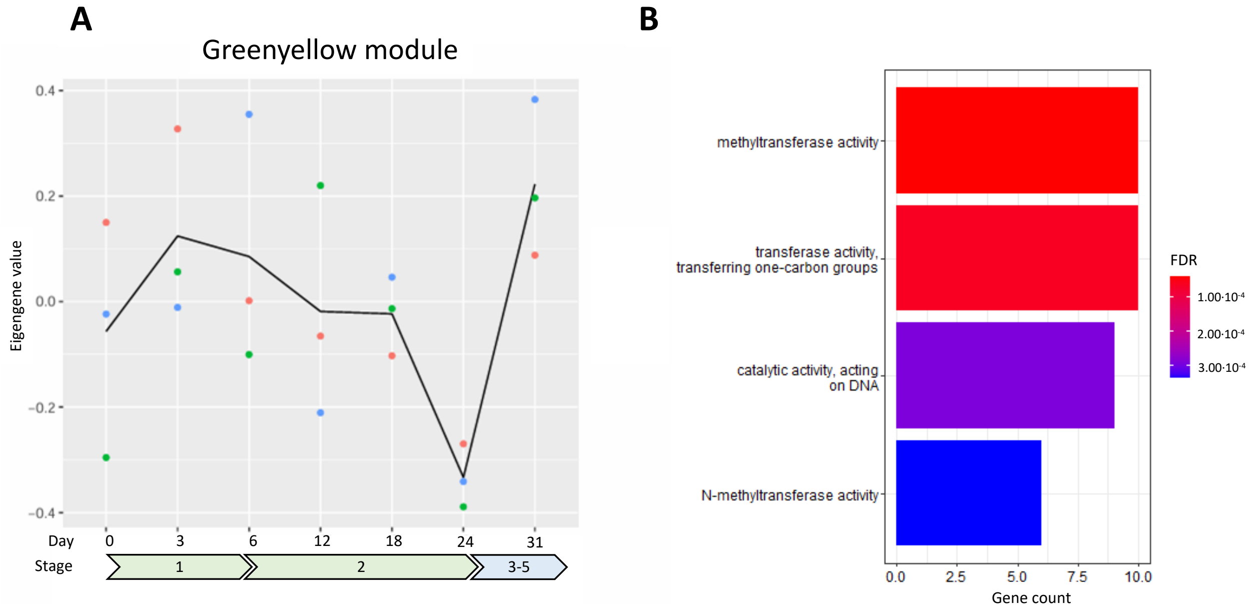


**Fig. S11** SC-IU green-yellow module results. **A)** Module eigengene are plotted over the course of hiPSC to SC differentiation. **B)** Functional enrichment analysis reveals significantly enriched ontology terms.

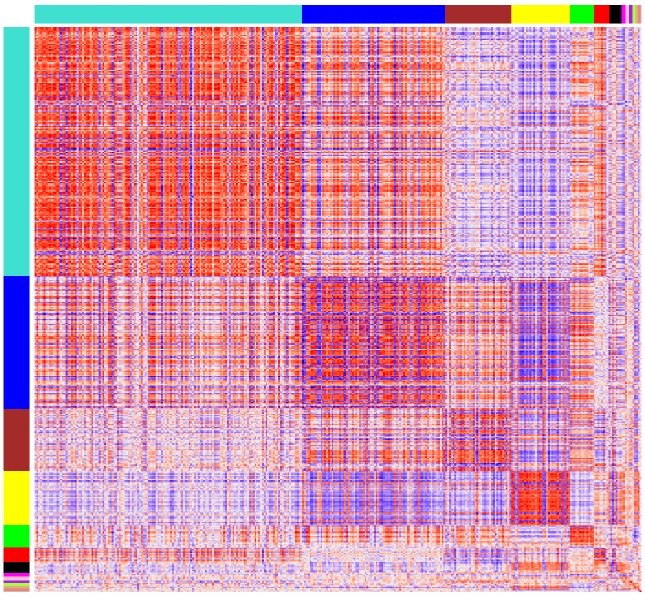

**Fig. S12** Heatmap of gene correlations from WGCNA analysis of the SNAT dataset . Gene co-expression modules are shown in the color-coded annotation tracks to the left and above the heatmap.

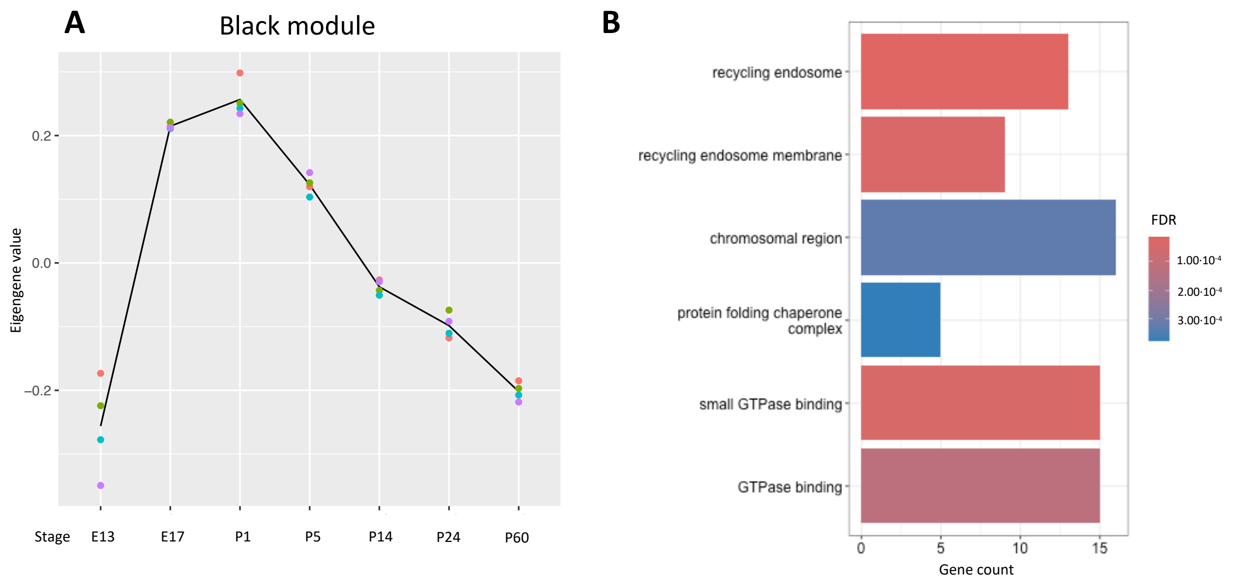


**Fig. S13** SNAT black module results. **A)** Module eigengene are plotted over the course of developmental stages. **B)** Functional enrichment analysis reveals significantly enriched ontology terms.

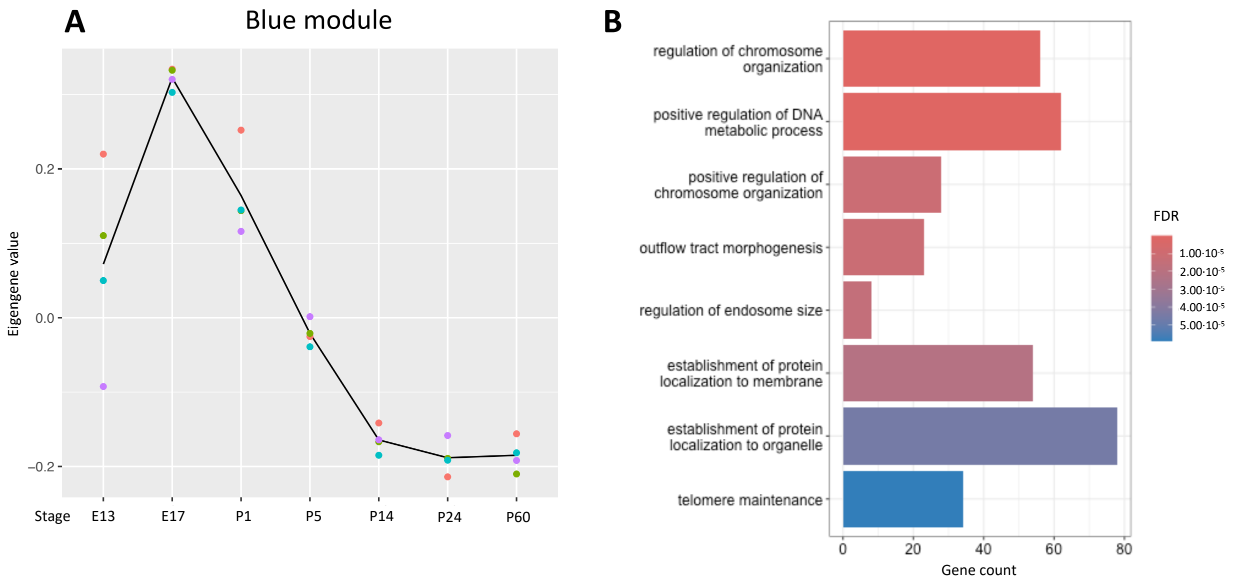


**Fig. S14** SNAT blue module results. **A)** Module eigengene are plotted over the course of developmental stages. **B)** Functional enrichment analysis reveals significantly enriched ontology terms.

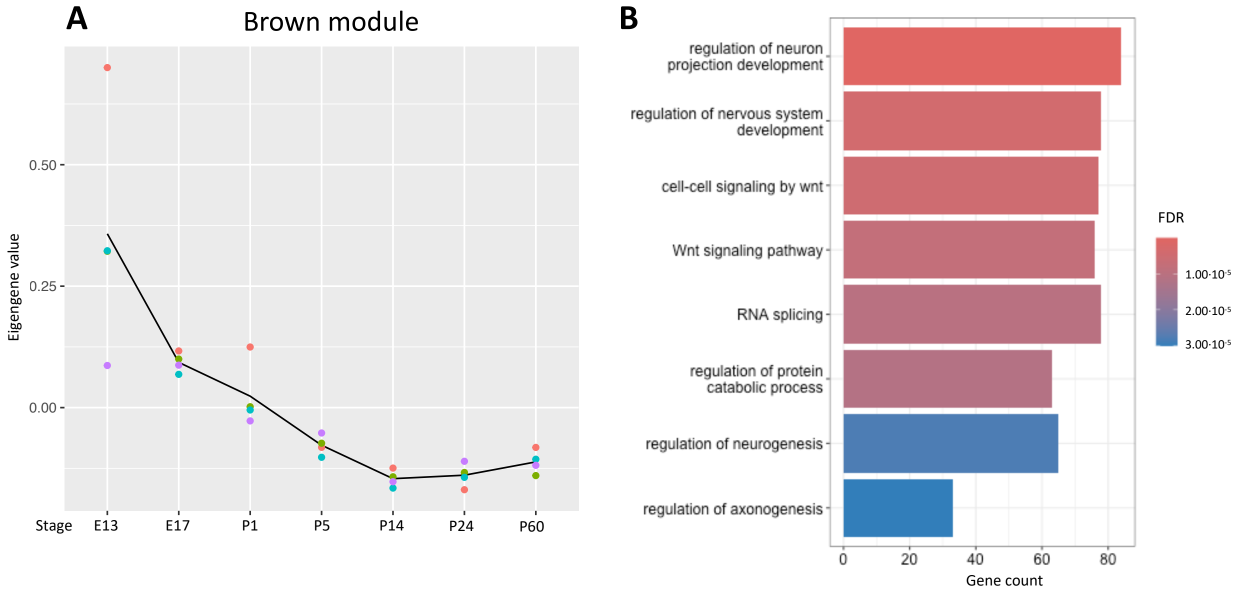


**Fig. S15** SNAT brown module results. **A)** Module eigengene are plotted over the course of developmental stages. **B)** Functional enrichment analysis reveals significantly enriched ontology terms.

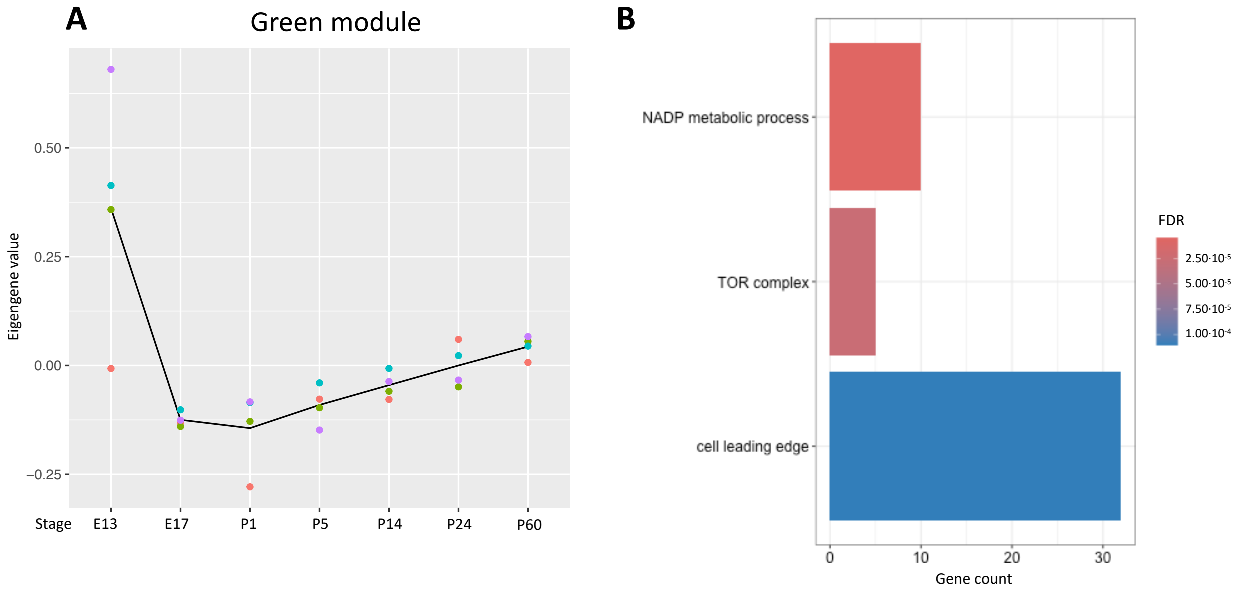


**Fig. S16** SNAT green module results. **A)** Module eigengene are plotted over the course of developmental stages. **B)** Functional enrichment analysis reveals significantly enriched ontology terms.

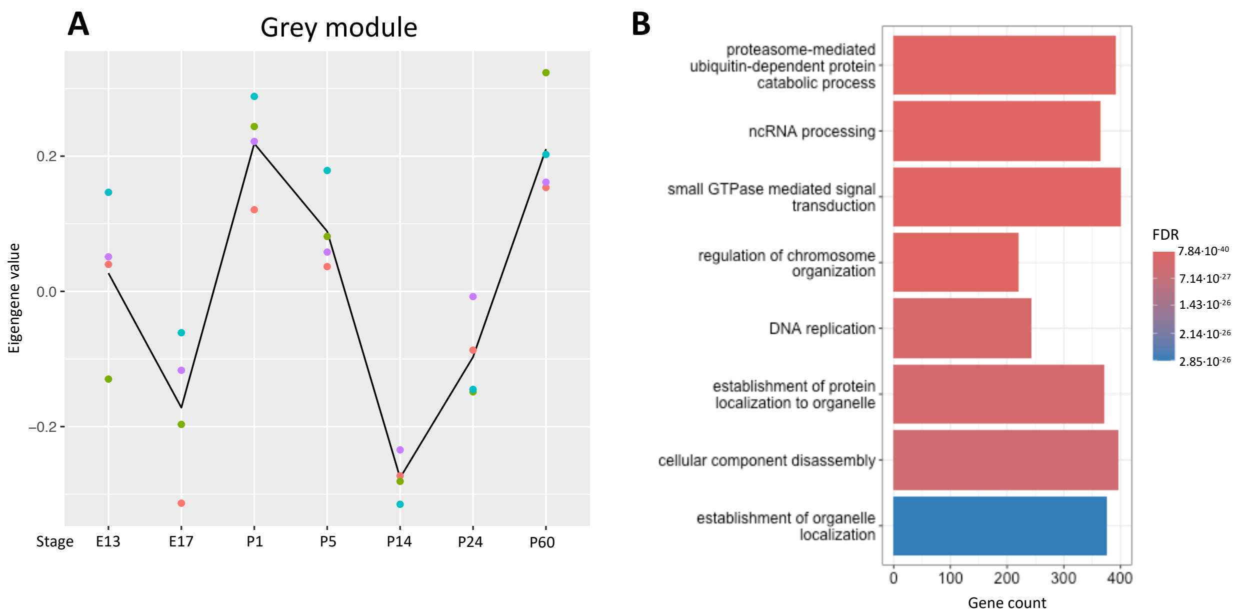


**Fig. S17** SNAT grey module results. **A)** Module eigengene are plotted over the course of developmental stages. **B)** Functional enrichment analysis reveals significantly enriched ontology terms.

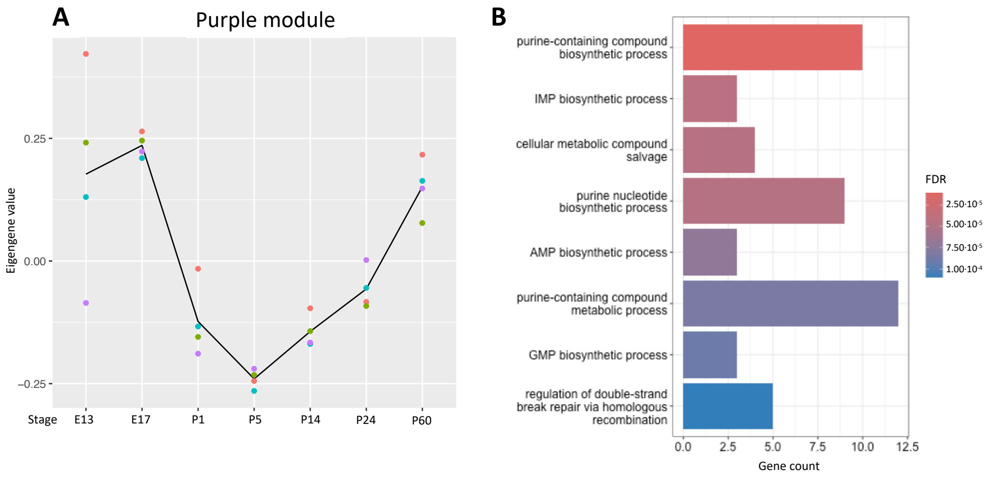


**Fig. S18** SNAT purple module results. **A)** Module eigengene are plotted over the course of developmental stages. **B)** Functional enrichment analysis reveals significantly enriched ontology terms.

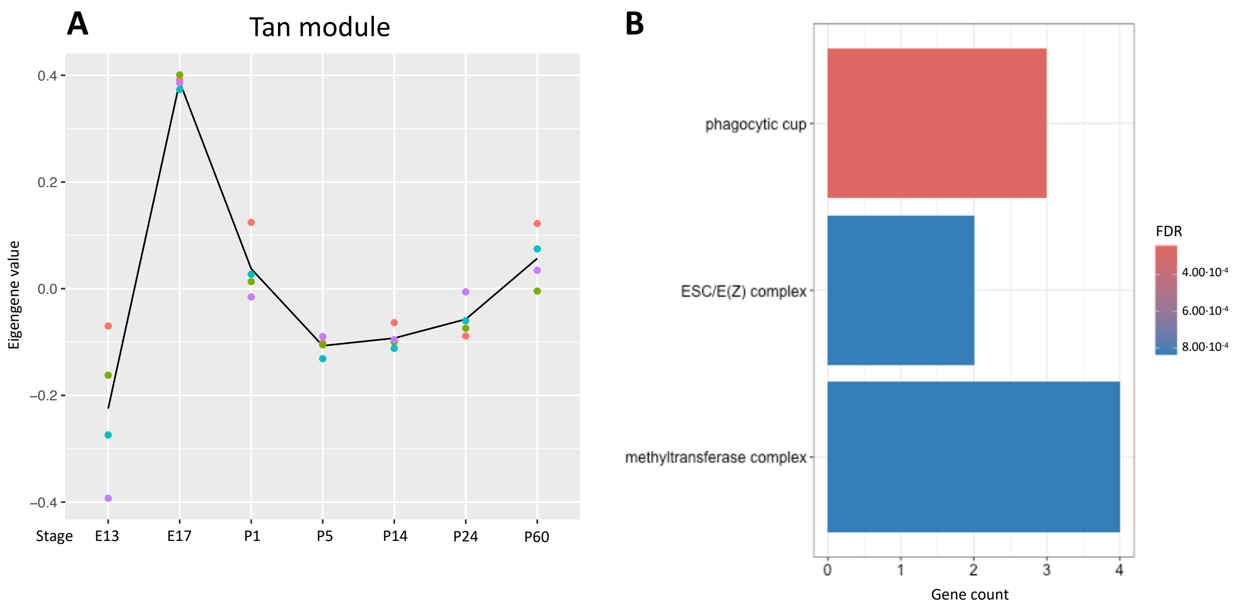


**Fig. S19** SNAT tan module results. **A)** Module eigengene are plotted over the course of developmental stages. **B)** Functional enrichment analysis reveals significantly enriched ontology terms.

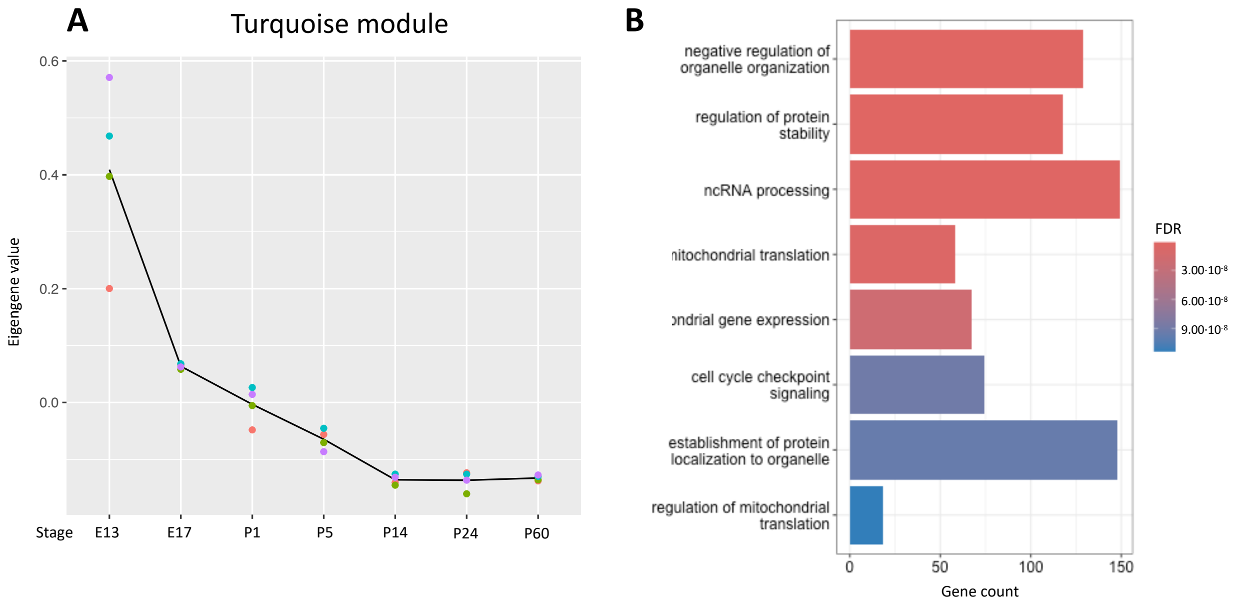


**Fig. S20** SNAT turquoise module results. **A)** Module eigengene are plotted over the course of developmental stages. **B)** Functional enrichment analysis reveals significantly enriched ontology terms.

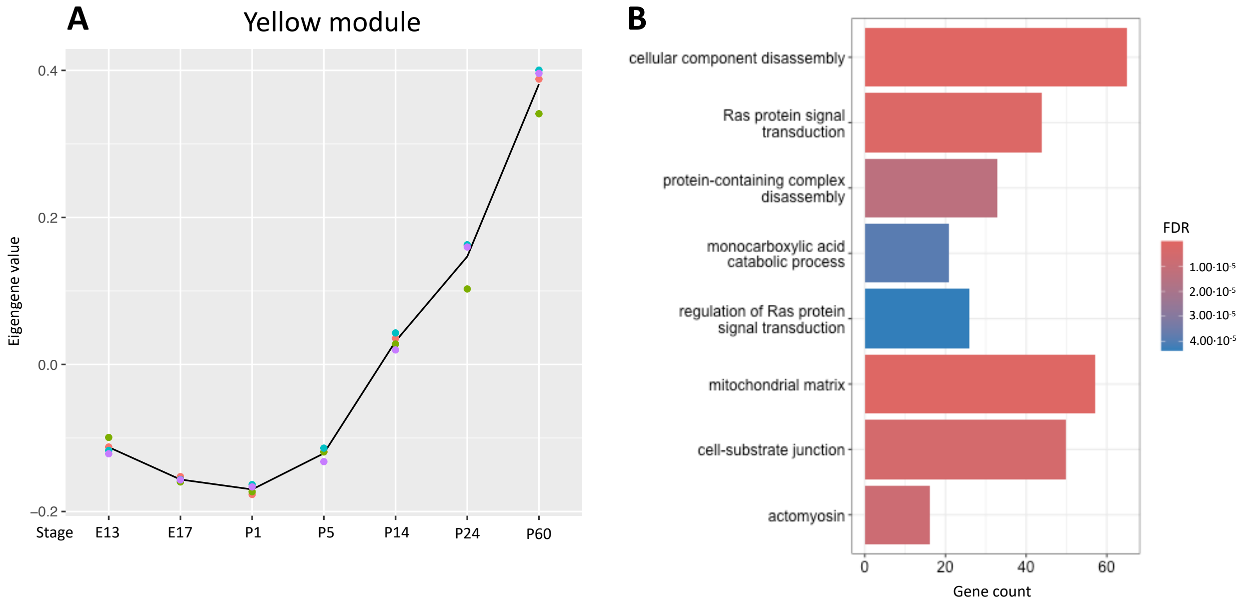


**Fig. S21** SNAT yellow module results. **A)** Module eigengene are plotted over the course of developmental stages. **B)** Functional enrichment analysis reveals significantly enriched ontology terms.

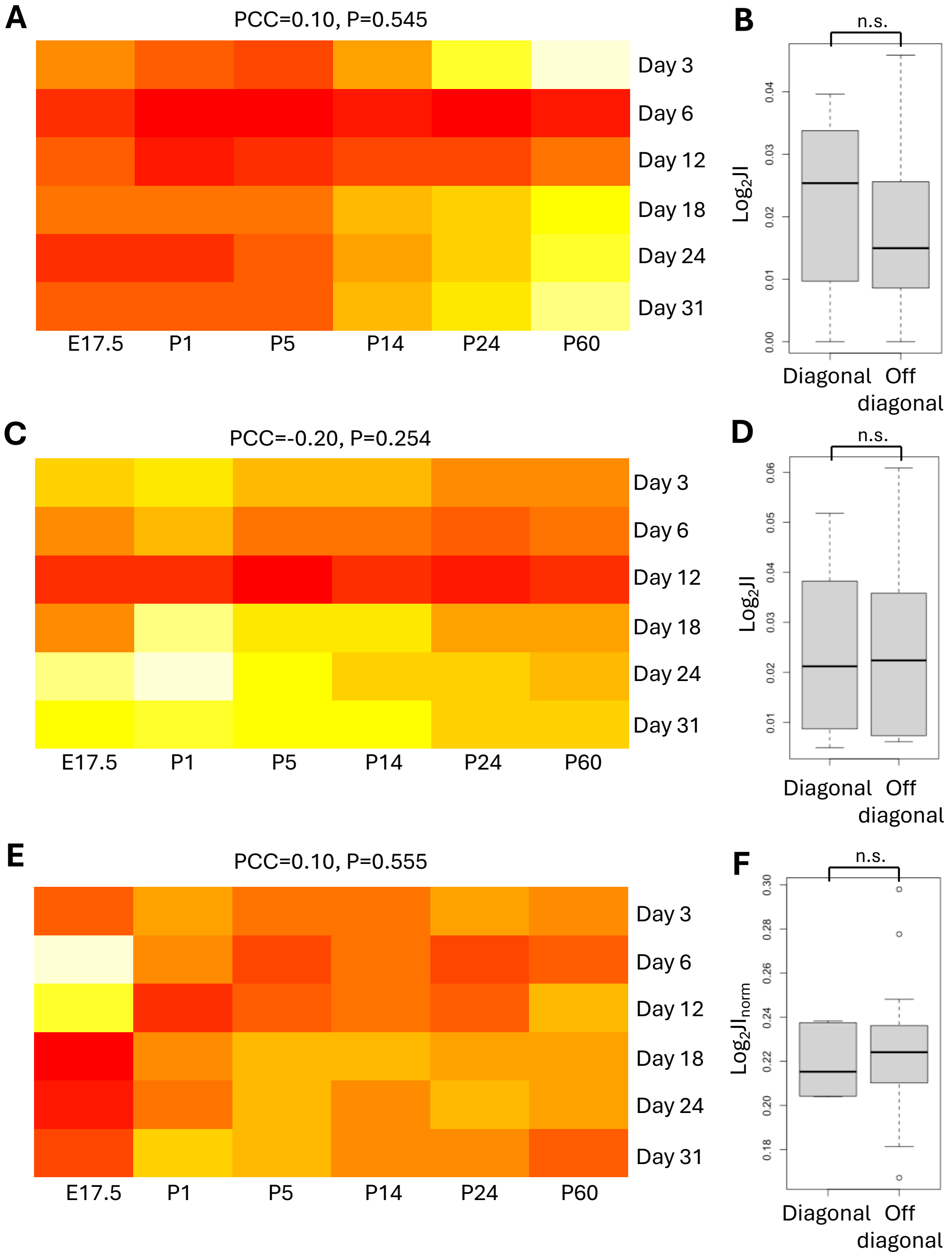


**Fig. S22** Correlation between uncorrected SC-IU and SNAT DEGs **A)** JI matrix of SC-IU differentiation up-regulated DEGs and SNAT up-regulated DEGs with comparison of JI matrix diagonal to non-diagonal **(B)**. **C)** JI matrix of SC-IU differentiation down-regulated DEGs and SNAT down-regulated DEGs with comparison of JI matrix diagonal to non-diagonal **(D)**. **E)** Normalized JI matrix of averaged SC-IU differentiation up/down-regulated DEGs and SNAT up/down-regulated DEGs with comparison of JI matrix diagonal to non-diagonal **(F)**.
